# Differences in expression of tumor suppressor, innate immune, inflammasome, and potassium/gap junction channel host genes significantly predict viral reservoir size during treated HIV infection

**DOI:** 10.1101/2023.01.10.523535

**Authors:** Ashok K. Dwivedi, David A. Siegel, Cassandra Thanh, Rebecca Hoh, Kristen S. Hobbs, Tony Pan, Erica A. Gibson, Jeffrey Martin, Frederick Hecht, Christopher Pilcher, Jeffrey Milush, Michael P. Busch, Mars Stone, Meei-Li Huang, Claire N. Levy, Pavitra Roychoudhury, Florian Hladik, Keith R. Jerome, Timothy J. Henrich, Steven G. Deeks, Sulggi A. Lee

## Abstract

The major barrier to an HIV cure is the persistence of infected cells that evade host immune surveillance despite effective antiretroviral therapy (ART). Most prior host genetic HIV studies have focused on identifying DNA polymorphisms (e.g., *CCR5Δ32*, MHC class I alleles) associated with viral load among untreated “elite controllers” (~1% of HIV+ individuals who are able to control virus without ART). However, there have been few studies evaluating host genetic predictors of viral control for the majority of people living with HIV (PLWH) on ART. We performed host RNA sequencing and HIV reservoir quantification (total DNA, unspliced RNA, intact DNA) from peripheral CD4+ T cells from 191 HIV+ ART-suppressed non-controllers. Multivariate models included covariates for timing of ART initiation, nadir CD4+ count, age, sex, and ancestry. Lower HIV total DNA (an estimate of the total reservoir) was associated with upregulation of tumor suppressor genes *NBL1* (q=0.012) and *P3H3* (q=0.012). Higher HIV unspliced RNA (an estimate of residual HIV transcription) was associated with downregulation of several host genes involving inflammasome (*IL1A, CSF3, TNFAIP5, TNFAIP6, TNFAIP9*, *CXCL3, CXCL10*) and innate immune (*TLR7*) signaling, as well as novel associations with potassium (*KCNJ2*) and gap junction (*GJB2*) channels, all q<0.05. Gene set enrichment analyses identified significant associations with TLR4/microbial translocation (q=0.006), IL-1β/NRLP3 inflammasome (q=0.008), and IL-10 (q=0.037) signaling. HIV intact DNA (an estimate of the “replication-competent” reservoir) demonstrated trends with thrombin degradation (*PLGLB1*) and glucose metabolism (*AGL*) genes, but data were (HIV intact DNA detected in only 42% of participants). Our findings demonstrate that among treated PLWH, that inflammation, innate immune responses, bacterial translocation, and tumor suppression/cell proliferation host signaling play a key role in the maintenance of the HIV reservoir during ART. Further data are needed to validate these findings, including functional genomic studies, and expanded epidemiologic studies in female, non-European cohorts.

**Author Summary:** Although lifelong HIV antiretroviral therapy (ART) suppresses virus, the major barrier to an HIV cure is the persistence of infected cells that evade host immune surveillance despite effective ART, “the HIV reservoir.” HIV eradication strategies have focused on eliminating residual virus to allow for HIV remission, but HIV cure trials to date have thus far failed to show a clinically meaningful reduction in the HIV reservoir. There is an urgent need for a better understanding of the host-viral dynamics during ART suppression to identify potential novel therapeutic targets for HIV cure. This is the first epidemiologic host gene expression study to demonstrate a significant link between HIV reservoir size and several well-known immunologic pathways (e.g., IL-1β, TLR7, TNF-α signaling pathways), as well as novel associations with potassium and gap junction channels (Kir2.1, connexin 26). Further data are needed to validate these findings, including functional genomic studies and expanded epidemiologic studies in female, non-European cohorts.

## Introduction

Despite several unique cases of possible HIV remission [1–3], there is still no HIV vaccine or cure. The major barrier to a cure is the persistence of infected cells that evade host immune surveillance despite effective antiretroviral therapy (ART). Modern antiretroviral therapy (ART) has transformed HIV disease into a treatable chronic disease for individuals who have access to, and are able to maintain, viral suppression [4]. However, ART alone does not eliminate persistent virus in most individuals [5, 6]. HIV cure trials aimed at reactivating and eliminating the HIV reservoir have thus far failed to show a clinically meaningful reduction in the HIV reservoir [7–12]. There is an urgent need to bridge drug discovery with a deeper understanding of host-viral dynamics. Although several host factors have been shown to influence the size of the “HIV reservoir”, such as the timing of ART initiation after initial HIV infection [13–16], maximum pre-ART viral load [17], ethnicity [17], and sex [17–20], there are few published human genomic and transcriptomic epidemiologic studies describing potential host factors influencing HIV persistence during treated infection.

Prior host genome wide association studies (GWAS) have focused on predictors of viral control (during untreated HIV disease), identifying key mutations in the C-C chemokine receptor type 5 gene (*CCR5Δ32*) and the human Major Histocompatibility Complex (MHC) human leukocyte antigen (HLA)-B and -C regions, that influence viral setpoint [21–24]. Recently our group reported these mutations (*CCR5Δ32 and HLA -B*57:01)* are associated with smaller HIV reservoir size [25]. However, mRNA expression of DNA variation is complex; the basal and/or the conditional expression of these genes in multicellular organisms are influenced by external controls (alternative splicing, polyadenylation, regulatory enhancers, etc.) which may differ by cell type and tissue [26–28]. The limited number of host gene expression studies during HIV infection (e.g., RNA sequencing) have compared gene expression between distinct clinical HIV groups. For example, one prior study compared gene expression among HIV “controllers” (individuals able to control virus in the absence of therapy) versus “non-controllers” [29]. Another study compared HIV non-controllers initiating ART “early” (<6 months from HIV infection) versus “later” (≥6 months after infection) [30]. However, no epidemiologic study has examined quantitative measures of the HIV reservoir size in relation to differences in host gene expression.

Here, we performed a cross-sectional study of 191 ART-suppressed HIV+ non-controllers to identify differentially expressed host genes in relation to three measures of the peripheral CD4+ T cell reservoir: HIV cell-associated “intact” DNA (an estimate of the frequency of potentially “replication-competent” virus with intact HIV genomes) [31], as well as total DNA (approximates intact + defective HIV DNA) and unspliced (full length transcript prior to alternative splicing) RNA. Increased expression of two putative tumor suppressor genes, *NBL1* and *P3H3*, was associated with smaller total HIV reservoir size (tDNA). Higher HIV usRNA was associated with downregulation of 17 host genes, including genes involved in pathogen pattern recognition (*TLR7*), inflammasome cytokine activation (*IL1A, CSF3, TNFAIP65, TNFAIP6, TNFAIP9*), and chemokine production (*CXCL3, CXCL10*). Higher usRNA also demonstrated a novel association with *KCNJ2*, a gene encoding for an inwardly rectifying potassium (Kir2.1) channel which has been shown to enhance HIV entry and release into host cells [32], as well as with *GJB2*, which encodes for a gap junction channel which facilitates cell-cell signaling (e.g., K+, Ca+, ATP) that has been implicated in cell-cell HIV transfer [33, 34]. These data add to the limited literature on host genetic predictors of the HIV reservoir and suggest that checks on cell proliferation might limit the total HIV reservoir size while a more “active” reservoir may stimulate host innate immune responses and inflammation during treated HIV disease. Further data are needed to validate these findings, including functional genomic studies using CRISPR-cas9 editing and longitudinal samples allowing causal inferences, as well as expanded studies in female, non-European cohorts.

## Results

### Study population

HIV+ ART-suppressed non-controllers were sampled from the UCSF SCOPE and Options cohorts (**Supplemental Fig 1**). The final 191 study participants were mostly male (96%) with a median age of 47 years and included individuals treated during early (within 6 months) or more chronic (>6 months after) HIV infection (**Table 1**). At the time of biospecimen collection, participants were ART-suppressed for a median of 5.1 years with a median nadir CD4+ T cell count of 352 cells/mm^3^ and maximum pre-ART HIV RNA of 5.1 log_10_copies/mL. As expected, our U.S.-based study population was diverse (**Fig 1)**. Thus, all results are shown for the total study population (adjusted for ancestry using principal components [35]), as well as restricted to the largest homogenous ancestral subgroup (Europeans), in order to enhance the ability to detect statistically significant genetic associations.

**Table 1.**
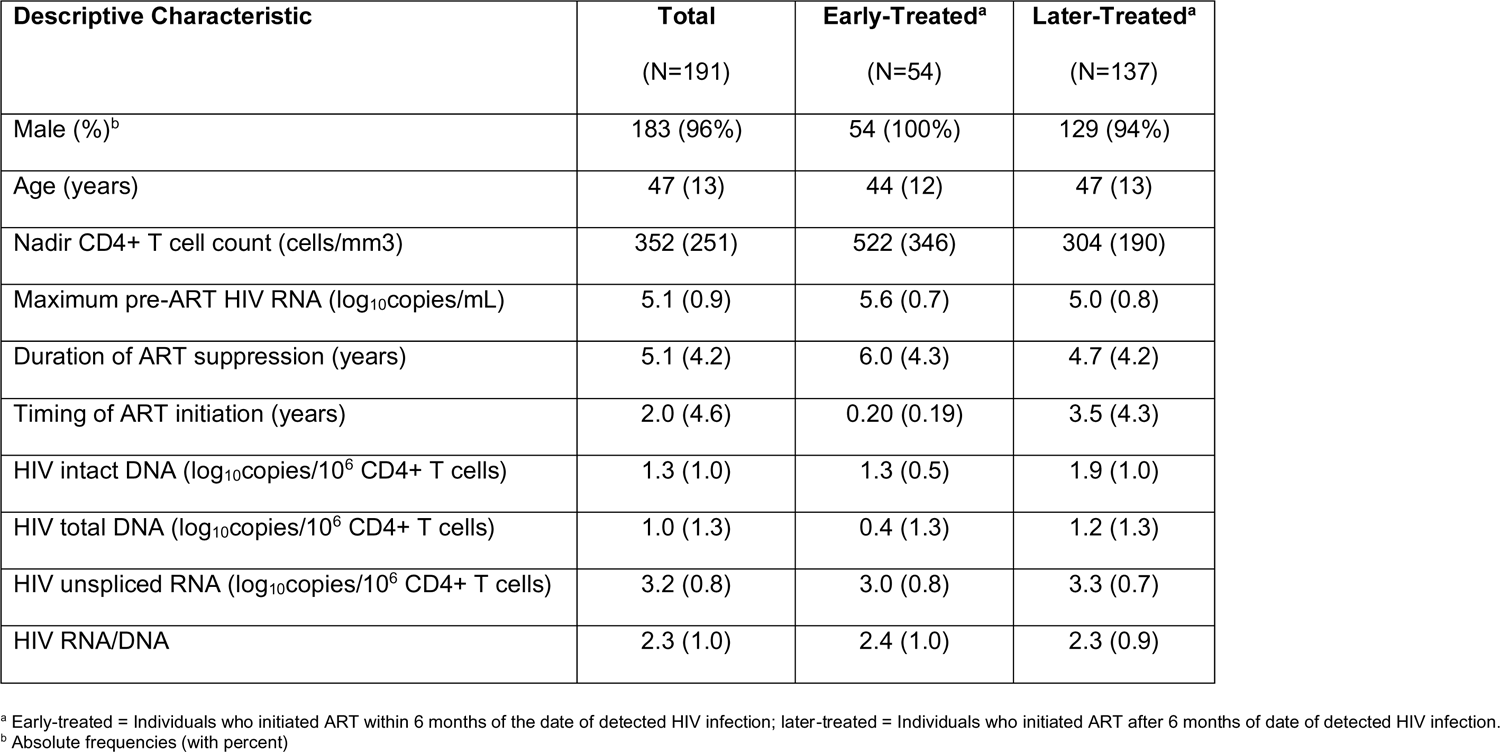
Descriptive statistics for the study population of 191 HIV-infected ART-suppressed non-controllers. Median frequencies (with interquartile ranges) are shown below unless otherwise specified.

**Figure 1.**
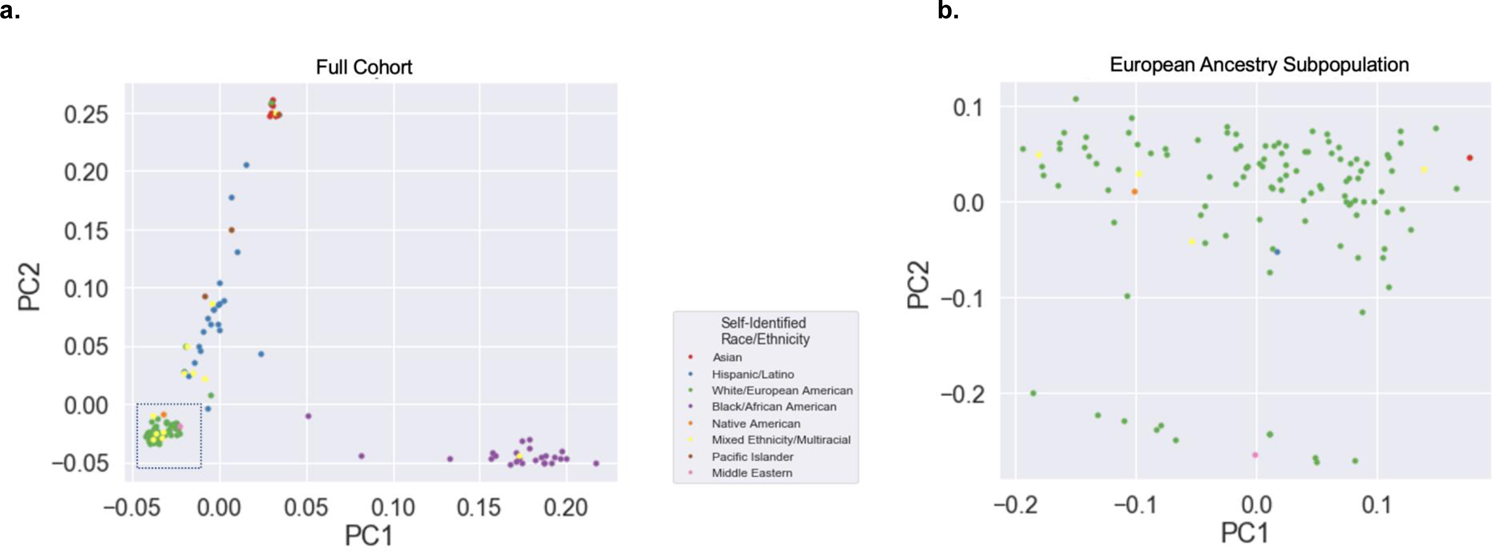
Principal component analysis (PCA) plot of the population structure. Principal component analysis (PCA) plot of the population structure of the full study cohort (a). Secondary PCA plot of the European ancestry subpopulation only (b) defined by the dashed box in the lower left of panel (a). Genetic PCs were calculated from genetic data from our whole exome analysis [25]. Most of the population was of European ancestry (bottom left of) (a) some continued variability. Some continued variability was observed in European ancestry subgroup (b). Self-identified race/ethnicity shown in the legend. Frequencies for participants were recorded as: White/European American (62%), Black/African American (14%), Hispanic/Latino (11%), Mixed Ethnicity/Multiracial (6%), Asian (4%), Pacific Islander (2%), Native American (<1%), and Middle Eastern (<1%).

### Measures of the HIV reservoir size were correlated with each other

Most of the HIV reservoir consists of cells harboring defective virus – i.e., cells that harbor HIV that is unable to go on to produce virions [36, 37], and yet the “replication-competent” reservoir is a major target of HIV eradication strategies [31, 38, 39]. Thus, there is currently no “gold standard” for measuring the HIV reservoir [40, 41]. Here, we performed three measures of the HIV reservoir from peripheral CD4+ T cells: total DNA (tDNA), unspliced RNA (usRNA), and HIV intact DNA. To estimate the frequency of the “replication-competent” reservoir, we performed a multiplexed droplet digital PCR (ddPCR) assay to quantify the frequency of cells with “intact” HIV sequences (i.e., likely to generate transcripts leading to virion production) [31, 40, 42]. HIV “total” (i.e., defective+intact) DNA and “unspliced” RNA (full-length HIV RNA) were also quantified using a separate, in-house quantitative polymerase chain reaction (qPCR) TaqMan assay [43]. HIV usRNA was statistically significantly correlated with tDNA (R=0.58, P=4.8×10^−19^) and intact DNA (R=0.24, P=1.9×10^−3^). However, HIV intact DNA was undetectable in 48% of our measured samples while total DNA was measurable in 95% of samples, which may have influenced the lack of association between tDNA and intact DNA in our study population (**Fig 2**).

**Figure 2.**
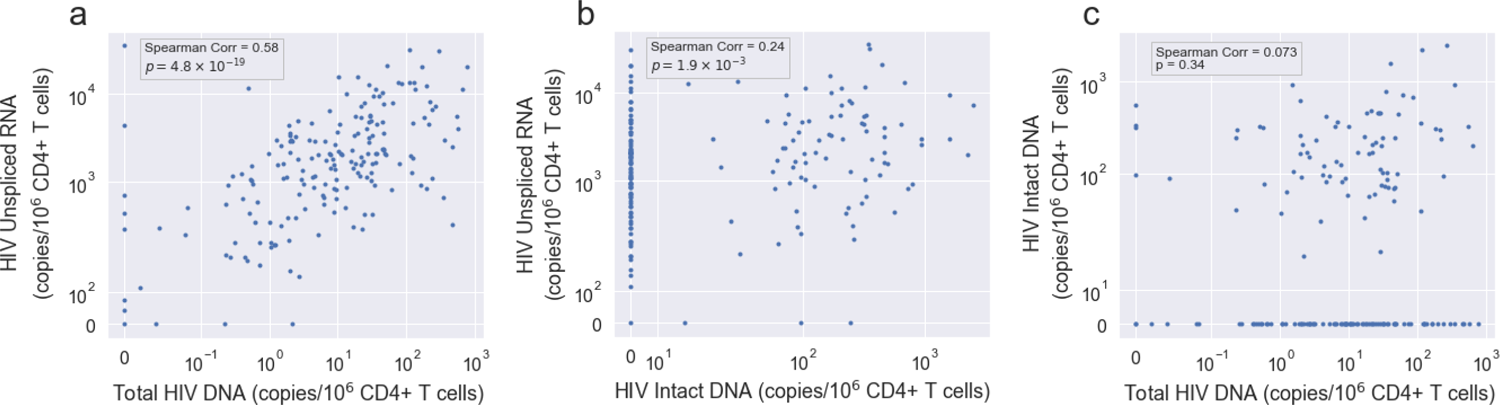
Correlations between three measures of HIV reservoir size. HIV unspliced RNA (usRNA) was significantly correlated with (a) HIV Total DNA (tDNA) and (b) HIV intact DNA; (c) tDNA and intact DNA were not correlated with one another.

### Earlier ART initiation and nadir CD4+ T cell count were associated with HIV reservoir size

Consistent with prior work [17, 37, 41], our study found that clinical factors previously shown to influence the size of the HIV reservoir were significantly associated with HIV reservoir measures quantified in our cohort. Earlier timing of ART initiation (<6 months from infection) was statistically significantly associated with lower levels of HIV intact DNA (R=0.21; p=6.5×10^−3^), tDNA (R=0.26; p=3.3×10^−4^), and usRNA (R=0.29; p=7.0×10^−5^) (**Fig 3**). Nadir CD4+ T cell count was associated with larger total HIV DNA reservoir size (R=−0.28; P=6.9×10^−5^), as well as higher levels of HIV usRNA (R=−0.28; p=9.6×10^−5^) and HIV intact DNA (R=−0.23; p=0.002). We did not observe a statistically significant association between HIV reservoir measures and other clinical factors: duration of ART suppression, age, or pre-ART HIV viral load, and we were unable to evaluate differences by sex/gender given low frequencies of female and transgender participants in our study.

**Figure 3.**
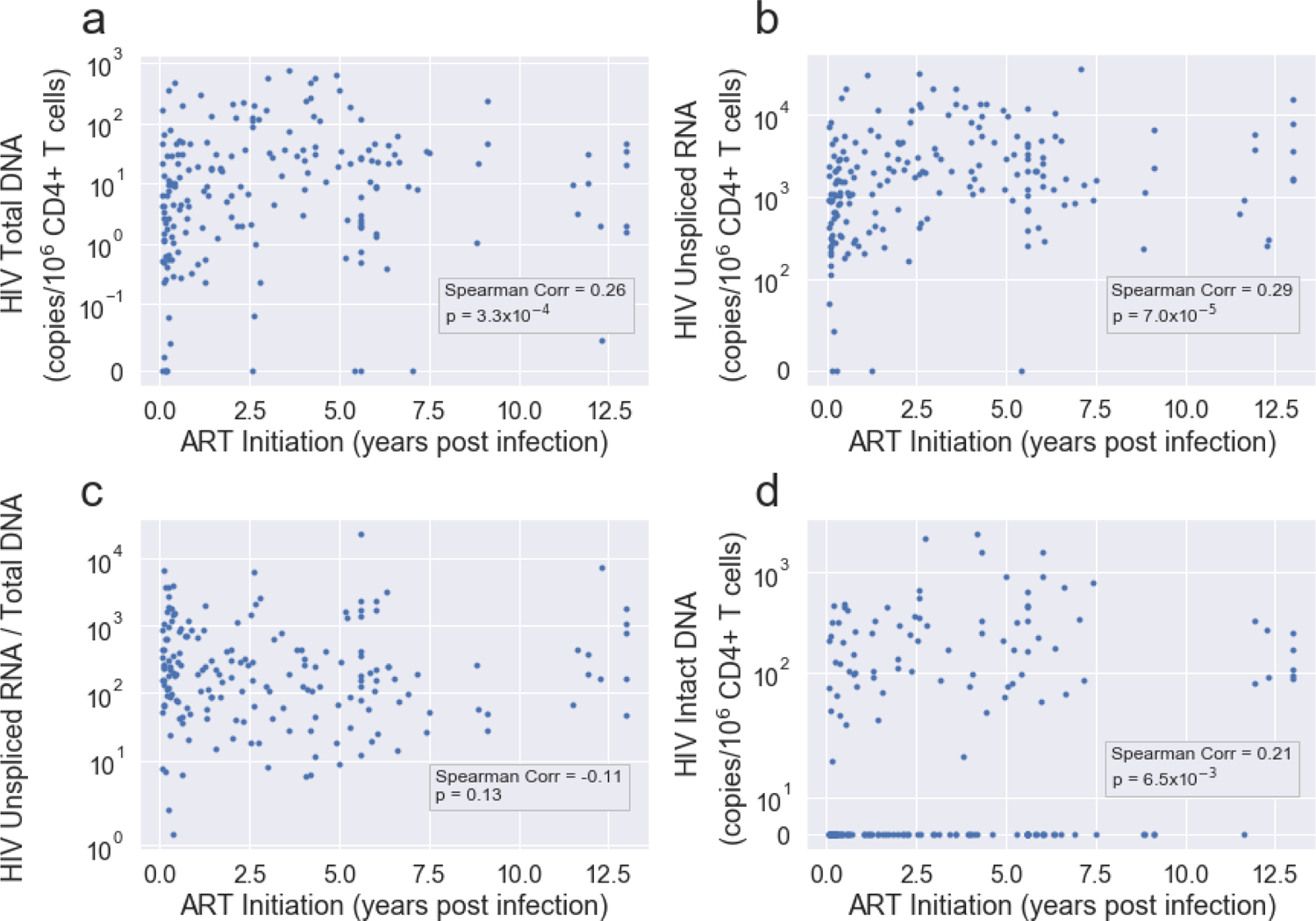
Measures of the HIV reservoir from peripheral CD4+ T cells were associated with timing of ART initiation. Panels A-D correspond to HIV tDNA, HIV usRNA, RNA/DNA, and Intact DNA, respectively. Spearman correlation and corresponding p-value are shown in each case. Earlier timing of ART initiation (<6 months from infection) was statistically significantly associated with smaller HIV intact DNA, tDNA, and usRNA.

### Increased expression of tumor suppressor genes was associated with total HIV DNA reservoir size while higher HIV usRNA was associated with downregulation of host inflammatory and innate immune genes

A total of 19,912 genes out of 60,719 were included for downstream differential gene expression analyses. In multivariate models adjusted for age, sex, nadir CD4+ T cell count, timing of ART initiation, ancestry (PCs), and residual variability (probabilistic estimation of expression residuals, PEERs), larger total HIV DNA reservoir size was statistically significantly associated with downregulation of two host tumor suppressor genes while higher HIV usRNA levels were associated decreased expression of 17 host genes involved in inflammation and innate immunity.

We observed that upregulation of tumor suppressor genes, *NBL1* and *P3H3, was* associated with smaller total HIV DNA reservoir size (**Supplemental Table 1**). For each fold-increase in gene expression of *NBL1* or *P3H3*, there was a statistically significant decrease in HIV total DNA (*NBL1*: −1.8%, q=0.012; *P3H3*: −1.6%, q=0.012). However, we observed the strongest associations between HIV reservoir size and host gene expression were with HIV unspliced RNA, largely reflecting the “transcriptionally active” HIV reservoir [44, 45]. A total of 17 host genes were inversely associated with HIV usRNA, including *KCNJ2* (−9.7%, q=0.003) which encodes for an inwardly rectifying potassium channel that has been shown to regulate HIV-1 entry and release [32], as well as *GJB2* (−7.1%, q=0.012), which encodes for a gap junction protein that facilitate cell-cell communication, potentially also cell-cell HIV transfer [46, 47]. In addition to these novel associations, HIV usRNA was also associated with several host genes involved in proinflammatory cytokine signaling and inflammasome activation (*IL1A*: −9.6%, q=0.012, *CSF3*: −7.5%, q=0.013; *TNFAIP6*: −7.6%, q=0.016, *TNFAIP9*: −6.9%, q=0.031, *TNFAIP5*: −5.9%, q=0.043), innate immune responses (*TLR7*: −7.1%, q=0.016), and chemokine production (*CXCL3*: −7.2%, q=0.043; *CXCL10*: −9.2%, q=0.049) (**Table 2, Supplement Table 2**). Given the large number of gene hits for HIV usRNA, we also performed network analyses to better visualize immunologic pathways identified from the differential gene expression analysis (q<0.25). We applied the ClueGo network analysis application, which clustered the large number of genes into biologically relevant, interpretable clusters [48]. These analyses highlighted several key pathways involving inflammasome activation [49–52] and bacterial translocation [53–55] – e.g., genes involved in NLRP3 (NOD-, LRR- and pyrin domain-containing protein 3) inflammasome activation, IL-1β, toll-like receptor 4, lipopolysaccharide (LPS), and IL-17 signaling (**Fig 4**). Using unbiased gene set enrichment analyses (GSEA), all genes in the transcriptome were rank-ordered by p-value to identify gene sets enriched for each HIV reservoir measurement. These analyses demonstrated that HIV total DNA was associated with complement activation and humoral immune response pathways, but these associations were only observed in the subgroup with the largest sample size, individuals of European ancestry (**Supplement Table 3**). HIV usRNA was again strongly associated with gene sets involving proinflammatory signaling and microbial translocation (“Response to Bacterium”, q=7.5×10^−5^; “Cellular Response to Lipopolysaccharide”, q=0.006), IL-1 signaling (“Interleukin-1 beta production”, q=0.008; “Regulation of Interleukin-1 Production”, q=0.008), and cytokine production (“Tumor Necrosis Factor Production”, q=0.006; “Tumor Necrosis Factor Superfamily Cytokine Production”, q=0.006; “Regulation of Tumor Necrosis Factor Production”, q=0.008). In addition, several gene sets related to IL-10 signaling (“regulation of interleukin-10 production”, q=0.037, “Interleukin-10 production”, q=0.041) – an anti-inflammatory pathway associated HIV immune dysregulation and persistence [56–58] – were also significantly associated with HIV usRNA (q=0.04) (**Fig 5, Supplement Table 4**).

**Table 2.** Differentially expressed host genes in relation to log_10_copies of HIV unspliced RNA (usRNA) in the full cohort (top panel) and the European ancestry subpopulation (bottom panel), at a Benjamini-Hochberg false discovery rate (FDR) of q<0.05. HIV usRNA was significantly associated with downregulation of 17 host genes, including *KCNJ2*, a novel association with a gene encoding for an inwardly rectifying potassium (Kir2.1) channel (which may enhance HIV entry and release into host cells [32]), gap junction (*GJB2*) as well as genes involved in pathogen pattern recognition (*TLR7*), inflammasome cytokine activation (*IL1A, CSF3, TNFAIP5, TNFAIP9, TNFAIP9*), and chemokine production (*CXCL3, CXCL10*). An additional list of host genes associated with HIV usRNA at an FDR q<0.25 is shown in **Supplemental Table 2**.

| HIV Unspliced RNA |  |  |  |  |  |  |
| --- | --- | --- | --- | --- | --- | --- |
| Gene | Gene Name | p<sup>a</sup> | q<sup>b</sup> | FC<sup>c</sup> | % Change<sup>d</sup> | Description |
| **Full Cohort** |  |  |  |  |  |  |
| *KCNJ2* | Potassium Inwardly Rectifying Channel Subfamily J Member 2, kir2.1 | 1.49E-07 | 0.003 | 0.903 | -9.7 | *KCNJ2*, encodes for an inwardly rectifying potassium channel (Kir2.1). Inwardly rectifying potassium ion channels can regulate HIV 1 entry and release into host cells [32]. Tight regulation of potassium ion concentrations has been shown to play a critical role in HIV-1 virus production in CD4+ T cells in cell culture models [143]. |
| *IL1A* | Interleukin-1 alpha | 1.55E-06 | 0.012 | 0.904 | -9.6 | *IL-1* is a potent proinflammatory cytokine that regulates inflammation by triggering a cascade of inflammatory mediators via NLRP3 inflammasome activation pathway [81, 82]. *IL-1* is an “upstream” pro-inflammatory inducer of interleukin-6 (*IL-6*) [83], which is the strongest biomarker for non-AIDS morbidity (e.g., myocardial infarction, stroke, malignancy) [84-87] and mortality [80, 86-89] among HIV-infected ART-suppressed individuals in resource-rich countries. |
| *GJB2* | Gap Junction Protein Beta 2 | 2.26E-06 | 0.012 | 0.929 | -7.1 | *GJB2* encodes for gap junction beta 2 protein, or connexin 26 (CX26), which acts as a communication channel between cells, facilitating transport of signaling molecules (calcium and cyclic AMP, ATP), and charged ions (K+, Ca+) [33, 34]. HIV-1 is thought to exploit these communication channels to disseminate infection and associated inflammation even in the absence of viral replication [46, 47]. |
| *KCNJ2-AS1* | KCNJ2 antisense RNA 1 | 2.49E-06 | 0.012 | 0.916 | -8.4 | *KCNJ2-AS1* is a long non-coding RNA (lncRNA) encodes for KCNJ2 antisense RNA 1. |
| *AC034199.1* | *AC034199.1* | 2.88E-06 | 0.012 | 0.946 | -5.4 | Novel transcript: no data found in literature in association with HIV or immune response. |
| *CSF3* | Colony Stimulating Factor 3 (G-CSF) | 3.93E-06 | 0.013 | 0.925 | -7.5 | *CSF3* encodes for granulocyte stimulating factor 3 (G-CSF), a member of the IL-6 superfamily of cytokines (78) and is also a growth factor for Neutrophils [90]. CSF3 modulates the function of CD4+ T cells by regulating cytokine production, their differentiation, and Treg induction [91]. |
| *TNFAIP6* | Tumor Necrosis Factor alpha induced protein 6 | 5.99E-06 | 0.016 | 0.924 | -7.6 | *TNFAIP6* encodes for TNF-stimulated gene 6 protein (TSG-6), which, like TNFAIP5 (TSG-5), is induced by tumor necrosis factor α (TNF-α) and interleukin-1 (IL-1) in response to lipopolysaccharide (LPS)-stimulation [101, 102]. |
| *TLR7* | Toll Like Receptor 7 | 6.59E-06 | 0.016 | 0.929 | -7.1 | *TLR7* is a member of toll-like receptor family of genes which plays critical role in pathogen recognition, activation of innate immune response and functions as a bridge between innate and adaptive immunity [127]. *TLR7*, is a pattern recognition receptor that can sense HIV single-stranded RNA (ssRNA) in endosomes [128, 129]. |
| *MRAS* | Muscle RAS oncogene homolog | 1.07E-05 | 0.024 | 0.945 | -5.5 | *MRAS* encodes for a protein in the Ras family of small GTPases which functions as signal transducers in cellular processes. |
| *TNFAIP9* | Tumor Necrosis Factor alpha induced protein 9 | 1.53E-05 | 0.031 | 0.931 | -6.9 | *TNFAIP9*, encodes for TNF-α induced protein 9 (TSG-9). It is also known as STEAP4 (six transmembrane epithelial antigen of prostate 4) involved in negative regulation of NF-κB, STAT-3 signaling, and IL-6 production [105, 106]. |
| *MIR3945HG* | MIR3945 Host Gene | 2.09E-05 | 0.038 | 0.942 | -5.8 | *MIR3945HG* is an interferon stimulated lncRNA. |
| *DAPK1-IT1* | DAPK1 Intronic Transcript 1 | 2.72E-05 | 0.043 | 0.950 | -5.0 | *DAPK1-IT1* is a lncRNA transcribed from the death associated protein kinases 1 (DAPK1). |
| *OR2B11* | Olfactory Receptor Family 2 Subfamily B Member 11 | 2.96E-05 | 0.043 | 0.939 | -6.1 | OR2B11 is a member of G-protein-coupled receptors (GPCR) responsible for the recognition and G protein-mediated transduction of odorant signals. |
| *CXCL3* | C-X-C Motif Chemokine Ligand 3 | 3.03E-05 | 0.043 | 0.928 | -7.2 | *CXCL3* is a member of CXC subfamily called cytokine-induced neutrophil chemoattractant (CINCs). *CXCL3* is involved in adhesion and migration of monocytes [109, 110], neutrophils chemoattraction [111-113], and angiogenesis [114]. |
| *TNFAIP5*<br>*/PTX3* | TNF Alpha-Induced Protein 5 (TNFAIP5), Pentraxin-related protein (PTX3), Tumor Necrosis Factor-Inducible Protein TSG-14 (TSG14). | 3.27E-05 | 0.043 | 0.941 | -5.9 | *TNFAIP5* is a pattern recognition receptor (PRRs) that is induced in response to TNF- $\alpha$, TLR engagement and IL-1β signaling [96, 97]. |
| *RRN3P4* | RRN3 Pseudogene 4 | 3.97E-05 | 0.049 | 0.950 | -5.0 | Pseudogene |
| *CXCL10* | C-X-C Motif Chemokine Ligand 10 | 4.21E-05 | 0.049 | 0.908 | -9.2 | *CXCL10* encodes for IP-10 (interferon gamma-induced protein 10) (interferon gamma-induced protein 10) which recruits activated Th1 lymphocytes to sites of infection [117-119] and in HIV, signals through TLR7/9-dependent pathways [119], predicts HIV disease progression [120, 121], correlates with acute HIV seroconversion [122], and promotes of HIV latency [123, 124]. |
| **European Ancestry Subpopulation** |  |  |  |  |  |  |
| *TLR7<sup>e</sup>* | Toll Like Receptor 7 | 1.48E-06 | 0.018 | 0.906 | -9.4 | *TLR7* is a member of toll-like receptor family of genes which plays critical role in pathogen recognition, activation of innate immune response and functions as a bridge between innate and adaptive immunity [127]. *TLR7*, is a pattern recognition receptor that can sense HIV single-stranded RNA (ssRNA) in endosomes [128, 129]. |
| *GJB2<sup>e</sup>* | Gap Junction Protein Beta 2 | 2.70E-06 | 0.018 | 0.909 | -9.1 | *GJB2* encodes for gap junction beta 2 protein, or connexin 26 (CX26), which acts as a communication channel between cells, facilitating transport of signaling molecules (calcium and cyclic AMP, ATP), and charged ions (K<sup>+</sup>, Ca<sup>+</sup>) [33, 34]. HIV-1 is thought to exploit these communication channels to disseminate infection and associated inflammation even in the absence of viral replication [46, 47]. |
| *AC034199.1<sup>e</sup>* | Ac034199.1 | 3.09E-06 | 0.018 | 0.930 | -7.0 | novel transcript |
| *PPP1R17* | Protein Phosphatase 1 Regulatory Subunit 17, | 4.20E-06 | 0.018 | 0.934 | -6.6 | PPP1R17 (Protein Phosphatase 1 Regulatory Subunit 17) is a substrate for cGMP-dependent protein kinase. |
| *IGSF6* | Immunoglobulin Superfamily Member 6 | 4.73E-06 | 0.018 | 0.972 | -2.8 | Immunoglobulin superfamily member 6 (IGSF6) is also known as downregulated by activation (DORA). |
| *AL133163.2* | AL133163.2 | 7.81E-06 | 0.025 | 0.945 | -5.5 | novel transcript |

<sup>a</sup> p = two sided p-value.
<sup>b</sup> q = two-sided false discovery rate (FDR) Benjamini-Hochberg q-value.
<sup>c</sup> FC = fold-change in host gene expression per two-fold change in copies of HIV from multivariate model adjusted for age, sex, nadir CD4+ T cell count, timing of ART initiation, ancestry (PCs), and residual variability (probabilistic estimation of expression residuals, PEERs).
<sup>d</sup> % Change = percent change in host gene expression per two-fold change in copies of HIV.
<sup>e</sup> % also significant in full cohort analysis.

**Figure 4.**
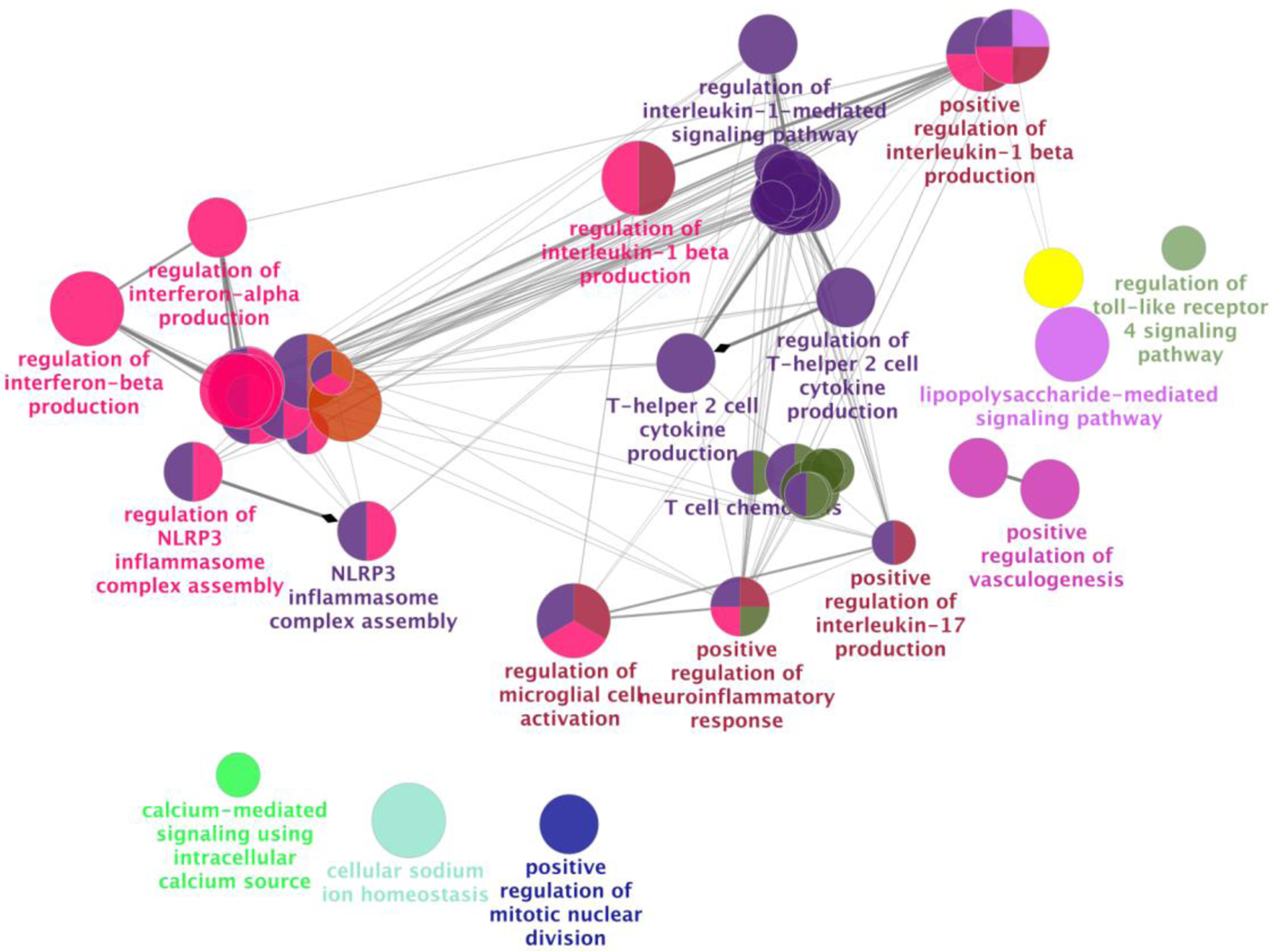
Network analysis of the top differentially expressed genes (see **Table 2** and **Supplemental Table 2**) associated with HIV unspliced RNA demonstrated that the top significant genes mapped to immunologic pathways involving bacterial translocation (e.g., TLR4 signaling, activated by bacterial lipopolysaccharide, LPS) and pro-inflammatory responses (e.g., IL-1β signaling, NLRP3 inflammasome assembly, Th2 cell cytokine production). A Benjamini-Hochberg false discovery rate (FDR) of q<0.05 was used to generate nodes (circles) based on kappa scores ≥0.4. The size of the nodes reflects the enrichment significance of the terms, and the different colors represent distinct functional groups.

**Figure 5.**
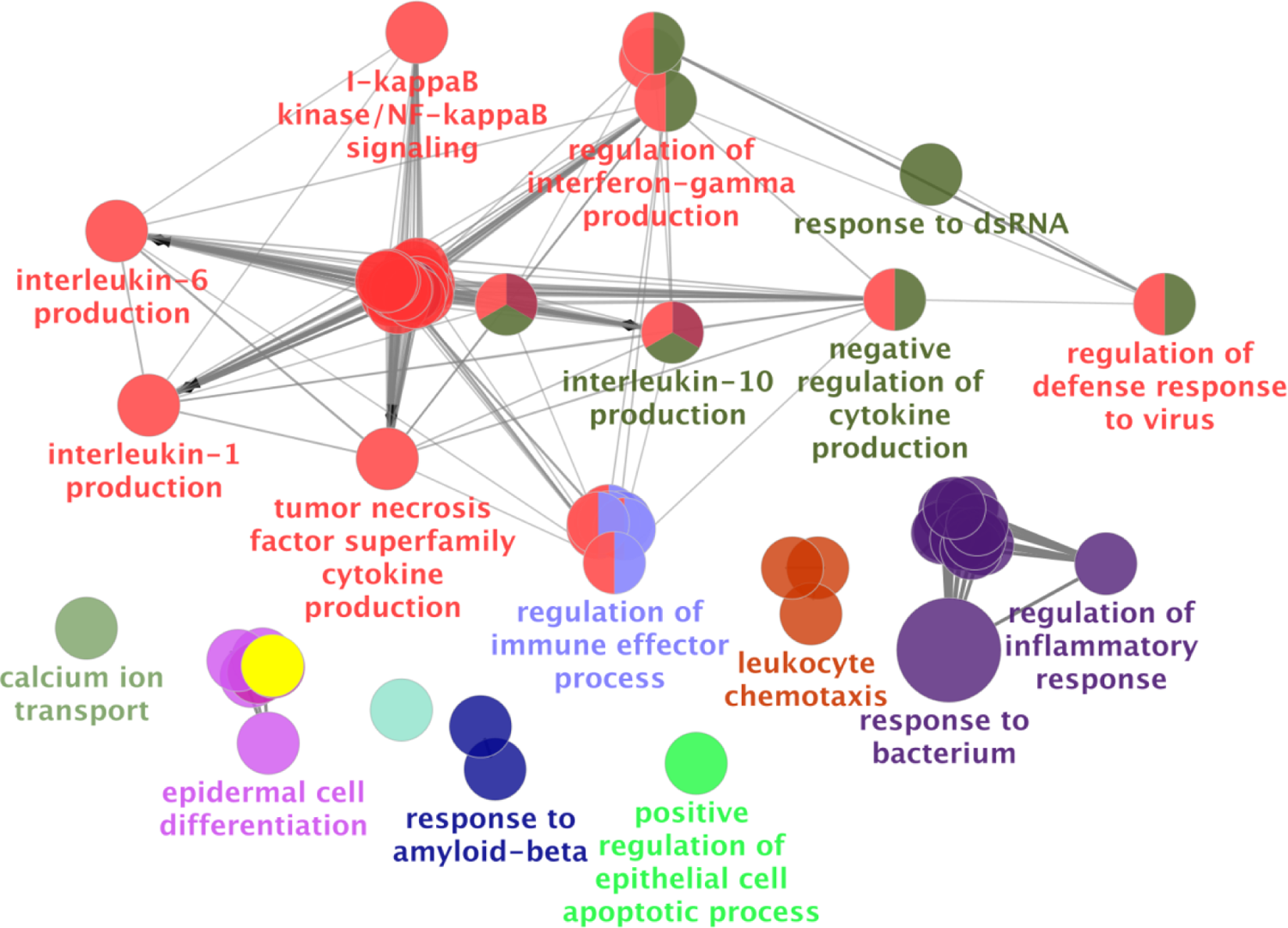
Network analysis of the top statistically significant gene sets associated with HIV unspliced RNA (see **Supplement Table 4**). Gene sets related to immunologic pathways involving bacterial translocation (e.g., response to bacterium, LPS-mediated signaling pathway), and inflammatory signaling (e.g., IL-1β], IL-6, IL-10, TNF-α), were significantly associated with HIV usRNA. A Benjamini-Hochberg false discovery rate (FDR) of q<0.05 was used to generate nodes (circles) based on kappa scores ≥0.4. The size of the nodes reflects the enrichment significance of the terms, and the different colors represent distinct functional groups.

### HIV intact DNA was undetectable in over half of the samples but was significantly associated with gene sets involving neutrophil activation in the European subgroup

HIV intact DNA was undetectable in over half of our measured samples while total DNA was measurable in 95% of samples (**Fig 2**). Hence, the statistical power to detect differential gene expression (DGE) associations was much lower for this assay compared to the other reservoir measures. By performing GSEA (a method that aggregates several genes into immunologically relevant “gene sets” to test for an association with HIV reservoir size), we were able to enhance the ability to detect potential associations with HIV intact DNA. In the differential gene expression analysis, among the European ancestry subgroup, we observed a positive trend (q<0.25) between HIV intact DNA and two genes, *PLGLB1* (+6.0%, q=0.23), which encodes for a protein that inhibits thrombus degradation, and *AGL* (+0.9%, q=0.23), encoding for an enzyme involved in glycogen degradation (**Supplemental Table 5**). GSEA demonstrated that gene sets involving neutrophil activation (“Neutrophil Degranulation”, q=0.046; “Neutrophil Activation Involved in Immune Response”, q=0.046; “Leukocyte Activation”; q=0.046) were significantly associated with HIV intact DNA, while gene sets reflecting myeloid-mediated immunity (“Myeloid Leukocyte Mediated Immunity”; q=0.058; “Myeloid Cell Activation Involved in Immune Response”; q=0.060) demonstrated a slight trend among European ancestry individuals (**Supplement Table 6**).

## Discussion

In the largest population-based transcriptomic HIV reservoir study to date, among HIV+ ART-suppressed non-controllers, we identified host genetic predictors (e.g., tumor suppressor genes) might act as “checks” on cell proliferation, potentially limiting the total HIV reservoir size. We also observed several associations with host genes indicating that a more “transcriptionally active” HIV reservoir [44, 45] may promote downregulation of potentially harmful host innate immune and proinflammatory responses. Given our cross-sectional study design, further functional and longitudinal epidemiologic studies are needed to determine potential causal relationships between host gene expression and HIV reservoir size. Nonetheless, these findings suggest that even during suppressive ART, ongoing host-pathogen dynamics maintain a delicate balance between a healthy host immune system and a persistent viral reservoir.

Differential gene expression analyses demonstrated that increased expression of tumor suppressor genes *NBL1* and *P3H3* was associated with a smaller total HIV DNA reservoir size. We observed statistically significant associations between two tumor suppressor genes, *NBL1* and *P3H3*, and HIV total DNA. Given the known function of these genes, these findings may suggest that increased expression of these host genes might impact the total HIV reservoir size, possibly by restricting cellular proliferation (**Supplemental Fig 2**). Alternatively, the association of these two tumor suppressor genes with total HIV reservoir size might suggest that cells harboring provirus integrated in genes promoting cell survival are selected for over time during ART suppression, as shown in HIV integration studies [59, 60]. *NBL1*, also known as neuroblastoma suppressor of tumorigenicity 1, is a transcription factor that belongs to the DAN (differential screening-selected gene aberrant in neuroblastoma) family of proteins [61, 62] and is involved in the negative regulation of cell cycle (G1/S transition) [63–66]. Interestingly, *NBL1* was differentially expressed in an RNA-seq *ex vivo* analysis of CD4+ T cells from rhesus macaques (after HIV-1 Env immunization and antibody co-administration) among groups that were treated with immune checkpoint modulators (CTLA-4, PD-1, and CTLA-4 + PD-1 Ab-treated), suggesting that NBL1 may be a potential pathway by which the cell cycle might be disrupted to enhance HIV-1 Env antibody responses [67]. *P3H3* encodes for Prolyl 3-Hydroxylase 3, which functions as a collagen prolyl hydroxylase (vital for collagen biosynthesis) that affects properties of the extracellular matrix and alters cellular behavior [68–71]. Prior studies suggest that *P3H3* plays a role as a tumor suppressor in breast, lymphoid, and other cancers [72–74]. Additional genes identified from the gene set enrichment analysis suggest that ongoing humoral immunity [75] and complement activation [76] may further contribute to maintaining the total HIV reservoir size during ART suppression (**Supplemental Table 3**).

HIV unspliced RNA was strongly associated with differential expression of several host genes previously associated with HIV disease, including genes involving inflammasome activation and inflammatory cytokine signaling (*IL1A, CSF3, TNFAIP5, TNFAIP6, TNFAIP9, IL10*), chemokine signaling (*CXCL3, CXCL10*), and innate immune response pathogen pattern recognition (*TLR7, TLR4*) (**Table 2, Supplemental Table 2, Figure 5, Supplement Table 5**). Given the known function of these genes and pathways, these findings might reflect that a more “transcriptionally active” HIV reservoir leads to host downregulation of potentially harmful proinflammatory signaling pathways during chronic treated HIV disease (**Supplemental Fig 3a**). This would be consistent with prior data demonstrating that excessive inflammation and immune activation predicts increased morbidity and mortality in HIV+ individuals despite effective ART [44, 45, 77–80].

Interleukin (IL)-1 is a potent proinflammatory cytokine that regulates inflammation, triggering a cascade of inflammatory mediators via the NOD-like receptor family pyrin domain containing 3 (NRLP3) inflammasome activation pathway [81, 82]. IL-1 is an “upstream” pro-inflammatory inducer of IL-6 [83], which is the strongest biomarker predicting serious non-AIDS morbidity (e.g., myocardial infarction, stroke, malignancy) [84–87] and mortality [80, 86–89] among HIV-infected ART-suppressed individuals. In our analysis, higher HIV usRNA was associated with decreased expression of *IL6* (−7.4%, q=0.062), *IL1A* (−9.6%, q=0.012), and *CSF3* (−7.5%, q=0.013) (**Table 2, Supplemental Table 2**); the latter encodes for granulocyte colony stimulating factor 3, G-CSF, a member of the IL-6 superfamily of cytokines [90] that modulates cytokine production, differentiation, and induction of Treg cells [91]. Our pathway-based analyses demonstrated a strong association between HIV usRNA and several genes in the IL-1β/NRLP3 inflammasome pathway (**Fig 4**), demonstrating for the first time, a link between this key immunologic pathway and the HIV reservoir.

Additional cytokines that were statistically significantly associated with HIV usRNA were several genes in the tumor necrosis (TNF)-α family: *TNFAIP5*, *TNFAIP6* and *TNFAIP9*. These genes encode for proteins regulating pro- and anti-inflammatory cellular signal transduction, differentiation, and apoptosis [92–95]. *TNFIAP5* encodes for a pattern recognition receptor that is induced in response to TNF-α, but also in response to toll-like receptor engagement and IL-1β signaling [96, 97], signals that are modulated through the NF-κB pathway [98]. *TNFAIP6* encodes for another TNF-α protein which functions as an anti-inflammatory protein [99, 100], is induced by IL-1 (upon LPS-stimulation) [101, 102], and interacts with *TNFAIP5* [103, 104]. *TNFAIP9*, also known as *STEAP4* (six transmembrane epithelial antigen of prostate 4), has been shown to negatively regulate NF-κB, STAT-3 signaling, and IL-6 production [105, 106]. Several chemokines were also inversely associated with HIV usRNA, again suggesting that a more “transcriptionally active” HIV reservoir might promote downregulation of host proinflammatory responses during long-term ART suppression. *CXCL3* and *CXCL10*, which encode for critical chemokines involved in the recruitment of neutrophils [107] and activated Th1 lymphocytes [108] to sites of inflammation respectively, were associated with a 7.2% and 9.2% decrease in gene expression per two-fold increase in HIV usRNA. CXCL3 regulates monocyte migration [109, 110], neutrophils chemoattraction [111–113], and angiogenesis [114], and is induced by proinflammatory IL-17 [115, 116]. *CXCL10* encodes for IP-10 (interferon gamma-induced protein 10) which recruits activated Th1 lymphocytes to sites of infection [117–119] and in HIV, signals through TLR7/9-dependent pathways [119], predicts HIV disease progression [120, 121], correlates with acute HIV seroconversion [122], and promotes HIV latency [123, 124]. Finally, in the gene set enrichment analysis, we also observed a statistically significant association between IL-10 signaling and HIV usRNA (**Fig 5, Supplement Table 4**). IL-10 is an immunosuppressive cytokine that plays an essential role in limiting the host immune response to pathogens and regulating the magnitude and duration of inflammation to prevent damage to the host [125]. IL-10 is broadly expressed by many immune cells, but cell type-specific signals also exist; IL-10 production is tightly regulated by changes in the chromatin structure, *IL10* gene transcription, and post-transcriptional regulatory mechanisms [126]. IL-10 has been associated with HIV immune dysregulation, e.g., impaired CD4+ T cell activation [58], and more recently, IL-10 has been shown to play a critical role in the maintenance of viral persistence [56, 57]. Among ART-suppressed PLWH, higher levels of IL-10 measured in blood and lymph nodes were significantly associated with HIV reservoir size (HIV integrated DNA) [57]. In SIV infected macaques, plasma IL-10 and IL-10 gene expression was associated with viral reservoir size (SIV DNA) in blood and lymph nodes, and *in vivo* neutralization of soluble IL-10 was shown to reduce B cell follicle maintenance [56].

There was also a statistically significant inverse association between HIV usRNA and genes associated with the host innate immune response (e.g., *TLR7*), while gene set enrichment also identified TLR4, associated with microbial translocation, to be significantly associated with HIV usRNA. These findings support the idea that HIV reservoir may not be entirely “quiescent” during ART and that ongoing residual viral transcription contributes to harmful persistent host immune activation even during ART suppression. *TLR7* encodes for a member of toll-like receptor family of genes which plays critical role in pathogen recognition, activation of the innate immune response, and functions as a bridge between innate and adaptive immunity [127]. TLR7 is a pattern recognition receptor that can sense HIV single-stranded RNA (ssRNA) [128, 129]. TLR7 agonist administration has been associated with delayed viral rebound [130] and reduced viral reservoirs in non-human primate studies [131]. A human clinical trial of the TLR7 agonist GS-9620 recently demonstrated a delay in viral rebound in HIV controllers after cessation of ART (NCT05281510) [132]. Interestingly, given that *TLR7* is located on the X chromosome, host *TLR7* transcriptional activity has been linked to acute viremia in HIV+ women (linked to type I interferon production) [133] as well as with enhanced innate immune function (i.e., plasmacytoid dendritic cell IFN-α and TNF-α production) [134]. Validation of our findings in female HIV+ cohorts will be critical for determining whether the host-viral dynamics described in our predominantly male study population are more pronounced in women, exhibiting a TLR7 signaling “dose-response” effect due to differential X inactivation in females [133]. Finally, another host innate pattern recognition receptor statistically significantly associated with HIV usRNA was *TLR4*, which was demonstrated in the pathway-based analyses linking *TLR4* to several gene sets involved in LPS-mediated signaling and IL-17 production (**Fig 4**). Our data add to prior studies linking bacterial gut translocation, systemic inflammation, immune activation, and HIV persistence [135–140].

The most statistically significant association with HIV usRNA was a novel association with *KCNJ2*, a gene that encodes for an inwardly rectifying potassium channel, Kir2.1. These potassium ion channels have been shown in prior lab studies to regulate HIV-1 entry and release [32]. In our analyses, a two-fold increase in HIV transcription (q=0.003) was associated with a 9.7% decrease in *KCNJ2* expression, as well as an 8.4% decrease in *KCNJ2-AS1* (encodes for KCNJ2 antisense RNA 1) expression (q=0.012). Potassium channels, including inwardly rectifying K+ channels, have been shown to regulate the life cycle of various viruses (e.g., Ebola [141], SIV [142]). Tight regulation of potassium ion concentrations have been shown to play a critical role in HIV-1 virus production in CD4+ T cells in cell culture models [143]. HIV Nef protein has been shown to increase K+ concentrations in cells [144], and in turn, changes in K+ concentration have been shown to regulate the HIV life cycle (e.g., viral entry, replication, and release) [32]. A small molecule inhibitor against Kir 2.1 has been recently identified [145]. The observed association between HIV usRNA and *KCNJ2*, as well with its antisense RNA, *KCNJ2-AS1*, might then suggest a potential novel mechanism – targeting specific types of potassium channels – to reduce the HIV reservoir size.

We also observed a novel association with *GJB2*, which encodes for gap junction beta 2 protein (also known as *CX26*, encoding for connexin 26). Gap junction proteins act as cell-cell communication channels to transport signaling molecules (e.g., K^+^, Ca+, ATP) [33, 34], and HIV-1 is thought to exploit these communication channels to disseminate infection as well as associated inflammation even in the absence of viral replication [46, 47]. As with *KCNJ2*, the observed association with *GJB2*, might suggest potential novel targets for limiting the HIV reservoir size.

We did not observe statistically significant associations with HIV intact DNA and host genes in the total study population. However, HIV intact DNA was undetectable in 48% of our measured samples, while for example, total DNA was measurable in 95% of samples (**Fig 2**). With so many samples below the limit of detection for intact DNA, the statistical power to detect differential gene expression is much lower for this assay than for the other HIV reservoir assays included in our study [146, 147]. Thus, we performed additional analyses restricted to the largest homogenous ancestral population (European ancestry subgroup) and performed pathway analyses to aggregate individual genes into immunologically relevant “gene sets” to test for an association with HIV reservoir size. In this way, we were able to enhance the ability to detect trends with HIV intact DNA. Higher HIV intact DNA was marginally associated with upregulation of *AGL* (involved in glycogen metabolism) [148–150] and *PLGLB1* (involved in thrombin clot degradation) [151–153] in the European ancestry subgroup (**Supplement Table 5**). Glycogen degradation involves breaking down stored glucose for immediate release and availability, and it has also been shown to play a key role in regulating the inflammatory immune response [150]. Antibody glycosylation has also been associated with inflammation-associated disease [150, 154], as well as time-to-viral rebound after ART interruption, a clinical definition of HIV reservoir size [148–150]. Besides its role in thrombolysis, *PLGLB1* has previously been associated with a replication-competent expanded HIV-1 clone described in a patient with squamous cell carcinoma (AMBI-1 integration) [155]; here it is associated with HIV intact DNA, which estimates the replication-competent HIV reservoir. Thus, the trends with *AGL* and *PLGLB1*, if further validated, might reflect that a larger “replication-competent” HIV reservoir contribute to vascular and metabolic complications that have been previously reported in HIV+ ART-suppressed individuals [44, 45, 77–80] (**Supplemental Fig 4**).

The study has several limitations that deserve mention. First, although the HIV reservoir has been shown to be relatively stable over time [17, 156, 157], our cross-sectional design provides a “snapshot” of the HIV reservoir after a median of 5.1 years of ART suppression and makes interpretation of the gene associations challenging. However, based on the known functions of the top gene hits, we conclude that some of the host genes identified in our analyses might reflect potential drivers of the HIV reservoir size (**Supplemental Fig 2**), while other host genes represent the impact of persistent HIV (**Supplemental Fig 3-4**). Indeed, the true *in vivo* associations might involve more complex feedback pathways between the HIV reservoir and host responses. Second, as is characteristic of our San Francisco-based HIV+ population, our study included mostly males of European ancestry. We accounted for this using well-established methods to adjust for population stratification bias [35, 158], as well as the use of PEERs, which help account for residual variance that often hampers RNA-seq data [159]. Nonetheless, it is important that these results be replicated in larger studies, especially those including women and individuals from different ethnic backgrounds. Third, the majority of the HIV reservoir persists in lymphoid tissues, not in the periphery [160]. However, recent data suggests that the tissue compartment largely reflects (and is the likely source of) the peripheral compartment [161–163]. Thus, it will be important to determine whether the results from our study are generalizable to the tissue HIV reservoir in future studies. Finally, we specifically chose to exclude HIV “elite” controllers in our study, since most people living with HIV do not fall within the ~1% of the HIV+ population able to suppress virus in the absence of therapy. Instead, the focus of our study was to determine other (uninvestigated) host gene expression associated with the HIV reservoir (signals that might be lost amidst a study population enriched for previously reported strong genetic effects, such as with HLA and/or *CCR5Δ32*).

Overall, our findings describe novel and immunologically relevant host genetic associations with the HIV+ reservoir. These include potential mechanisms inhibiting cell proliferation to limit the size of the overall HIV reservoir, as well as compensatory host downregulation of harmful persistent innate immune activation and inflammation (e.g., toll-like receptor, IL-1β/NRLP3 inflammasome, microbial translocation, IL-10 signaling etc.). Finally, the strongest association with HIV transcription was with *KCNJ2*, a potential novel mechanism by which the host restricts residual HIV propagation via inwardly rectifying potassium channels. Additional studies are needed to validate these findings using approaches functionally and epidemiologically like CRISPR-Cas9 editing and expanding these studies to include more diverse patient populations, including female and non-European ancestry individuals, using longitudinal samples.

## Materials and Methods

### Study Participants

HIV+ ART-suppressed non-controllers from the UCSF SCOPE and Options HIV+ cohorts were included in the study. Inclusion criteria were laboratory-confirmed HIV-1 infection, availability of cryopreserved peripheral blood mononuclear cells (PBMCs), and plasma HIV RNA levels below the limit of assay quantification for at least 24 months at the time of biospecimen collection. We excluded individuals HIV “elite controllers” to focus on genetic variants that drive HIV persistence among non-controllers during ART suppression but also analyzed previously reported strong genetic effects associated with HIV+ elite control [164–166]. The estimated date of detected infection (EDDI) was calculated for each study participant to determine recency of infection in relation to ART initiation using the Infection Dating Tool (https://tools.incidence-estimation.org/idt/) [167]. Additional exclusion criteria were potential factors that might influence HIV reservoir quantification, including recent hospitalization, infection requiring antibiotics, vaccination, or exposure to immunomodulatory drugs in the six months prior to sampling timepoint. The research was approved by the UCSF Committee on Human Research (CHR), and all participants provided written informed consent.

### HIV Reservoir Quantification

Cryopreserved PBMCs were enriched for CD4+ T cells (StemCell, Vancouver, Canada), and DNA and RNA were extracted from CD4+ T cells using the AllPrep Universal Kit (Qiagen, Hilden, Germany). Cell-associated total HIV DNA and unspliced RNA were quantified by an in-house quantitative polymerase chain reaction (qPCR) TaqMan assay using HIV-1 long terminal repeat (LTR)-specific primers as previously described [43]. Participant specimens were assayed with up to 800 ng of total cellular RNA or DNA in replicate reaction wells and copy number determined by extrapolation against a 7-point standard curve (1–10,000 copies/second) performed in triplicate. HIV intact DNA was quantified by targeting five regions on the HIV genome, including highly conserved regions and positions that are frequently deleted or hypermutated [31]. Optimized restriction enzyme digestion was used to prepare the genomic DNA for droplet formation while minimizing the amount of shearing within the viral genome. The protocol targeted 5 regions in the HIV genome across two droplet digital PCR (ddPCR) assays. Droplet generation and thermocycling were performed according to manufacturer instructions. This multiplex ddPCR assay allowed the analysis of potentially replication-competent (“intact”) proviral genomes by quantifying the number of droplets positive for 3 targets per assay. Two targets in a housekeeping gene (*RPP30*) were used to quantify all cells, and a target in the T cell receptor D gene (*TRD*) was used to normalize the HIV copy numbers per 1×10^6^ CD4+ T cells. A DNA shearing index (DSI) was then calculated, and mathematically corrected for residual DNA shearing as measured by *RPP30* targets to calculate the estimated number of intact proviral genomes per million CD4+ T cells after correcting for shearing [42].

### Host RNA sequencing

A separate aliquot of the extracted RNA from CD4+ T cells was then used to perform host RNA sequencing. HTStream pre-processing pipeline (s4hts.github.io/htstream/) was used for removing PCR duplicates, adapters, N characters, PolyA/T sequences, Phix contaminants, and poor-quality sequences (with quality score <20 with sliding window of 10 base pairs). The quality of raw reads was assessed using FastQC [168]. All samples had a per base quality score and sequence quality score >30. RNA-seq reads were then mapped to the human genome (GRCh38) [169] with a corresponding annotation file from the GENCODE project [170]. Alignment and gene quantification were performed using the STAR alignment tool and its quantification protocols [171–173]. Gene expression was converted to counts per million (CPM). To normalize the distribution of expression values across the experiment, the trimmed mean of M-values (TMM) [174] was used for sample-specific adjustment. Low-expressed genes (<1 CPM for all samples) were removed. The mean-variance trend was estimated [175] to assign observational weights based on predicted variance on log2-counts per million (log-CPM) using the Limma-Voom pipeline [176].

### Differential Gene Expression Analysis

Multivariate linear models were fit for each of the three measures of the HIV reservoir size using the Limma-Voom workflow [175, 176], a quantitative weighting method that utilizes variance modeling to accommodate for residual technical and/or biological heterogeneity [175]. For all analyses, in order to account for potential population stratification bias (i.e., systematic differences in results due to ancestry rather than association of genes with disease) we used well-established methods to account for this by (1) calculating and including the first five principal components (PCs) as covariates in the multivariate models [35] and (2) performing sensitivity analyses among the largest subgroup, individuals of European ancestry. Eigenvalues were calculated to generate genetic principal components (PC) to adjust for ancestry [35]. Multivariate models also included covariates for sex, age, timing of ART initiation, and nadir CD4+ T cell count (duration of ART suppression and maximum pre-ART viral load did not significantly improve the fit of the models and were not included as covariates in the final models), as well as PEERs (probabilistic estimation of expression residuals) to control for additional systematic sources of bias [159]. Model fit was assessed using a lambda genomic coefficient close to 1 [177]. Statistical significance was determined using a false discovery rate (FDR) q-value threshold of <0.05.

### Gene Set Enrichment Analyses and Network Analyses

For each of the three HIV reservoir measures, we also performed gene set enrichment analyses (GSEA) to more broadly evaluate whether specific immune pathways were linked to each HIV reservoir measurement. Genes from the entire transcriptome were first rank-ordered by q-values from the differential gene expression analysis for each HIV reservoir measure, and then the rank-ordering was used to identify immunologic pathways that were enriched from our dataset, using the Gene Ontology Biological Processes (GO-BP) database [178]. For the HIV usRNA analyses, for which there were several statistically significant differentially expressed genes (even after multiple-testing), we performed network analyses to better cluster and visualize the statistically significant results. Using ClueGo, a network analysis application [48], only statistically significant and marginally significant genes (q<0.25) were included to calculate Kappa statistics that allowed more meaningful visualization of potential biologically relevant pathways (**Fig 4**).

## Acknowledgements

The authors wish to acknowledge the participation of all the study participants who contributed to this work as well as the clinical research staff of the SCOPE and Options who made this research possible. All funders had no role in study design, data collection and analysis, decision to publish, or preparation of the manuscript. All authors provided critical feedback in finalizing the report. SAL and DAS conceived and designed the study with critical feedback from SGD, TJH, and DAS. SGD, JM, FH, CP, RH, and SAL coordinated the collection, management, and quality control processes for the cohort clinical data and SGD, JM, FH, CP, MPB, MS provided biospecimens. SAL, CT, JM, KB, TP, EAG performed participant sample processing, SAL performed the RNA sequencing assays, and AKD performed quality control analyses and the association analyses for the study under the guidance of SAL and DAS. SAL, CT, and KH performed the qPCR HIV reservoir assays (total DNA, unspliced RNA) in the lab of TH. CNL and MLH performed the ddPCR HIV reservoir assay (intact DNA) in the labs of FH, KRJ, and HPK. AKD, PR, DAS, TJH, and SAL analyzed these HIV reservoir data in relation to host transcriptomic and clinical phenotype data. AKD, SAL, and DAS wrote the report. All authors provided critical feedback in finalizing the manuscript.

