## Supplemental Files for "Differences in expression of tumor suppressor, innate immune, inflammasome, and potassium/gap junction channel host genes significantly predict viral reservoir size during treated HIV infection"

**Supplemental Figure 1.** Study participant sample selection flowchart. Specific inclusion and exclusion criteria are listed for each selection step and for HIV reservoir measure analysis.

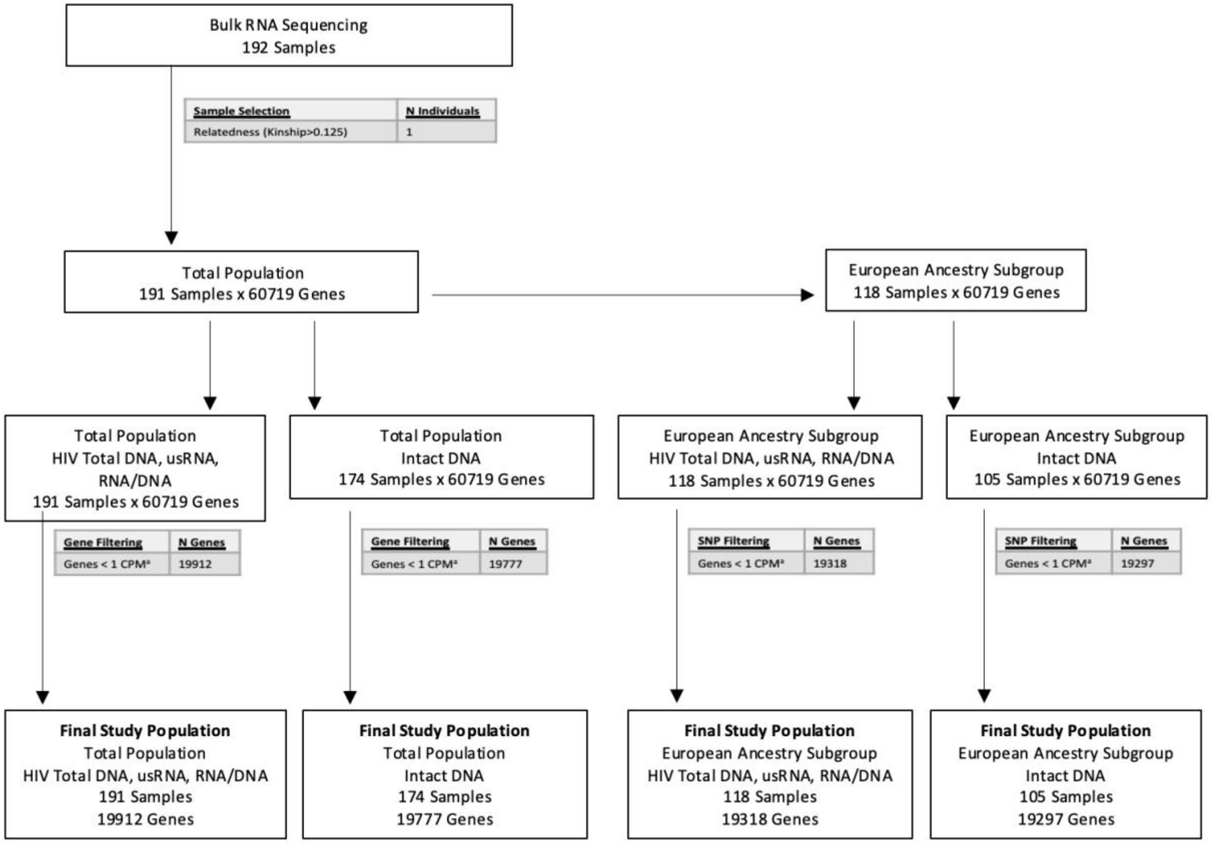


[a] p = Counts per million

**Supplemental Table 1.** Differentially expressed host genes associated with HIV total DNA (tDNA) in the total study population (top panel) and the European ancestry subgroup (bottom panel), at a Benjamini-Hochberg false discovery rate (FDR) of q<0.05.

| HIV Total DNA | | | | | | |
|---|---|---|---|---|---|---|
| **Gene** | **Gene Name** | **p[a]** | **q[b]** | **FC[c]** | **% Change[d]** | **Description** |
| **Total Study Population** | | | | | | |
| *NBL1* | NBL1, DAN Family BMP Antagonist | 6.14E-07 | **0.012** | 0.982 | -1.8 | *NBL1* is also known as neuroblastoma suppressor of tumorigenicity 1. NBL1 is a transcription factor that belongs to the DAN (differential screening-selected gene aberrant in neuroblastoma) family of proteins [61, 62] and negatively regulates cell cycle (G1/S transition) [63-66]. |
| *P3H3* | Prolyl 3-Hydroxylase 3 | 1.25E-06 | **0.012** | 0.984 | -1.6 | *P3H3*, functions as a collagen prolyl hydroxylase (vital for collagen biosynthesis), alters the cellular behavior by effecting properties of extracellular matrix [68-71], and studies suggest that it plays a role as a tumor suppressor in breast, lymphoid, and other cancers [72-74]. |
| **European Ancestry Subgroup** | | | | | | |
| NA | | | | | | |

[a] p = two sided p-value.
[b] q = two-sided false discovery rate (FDR) Benjamini-Hochberg q-value.
[c] FC = fold-change in host gene expression per two-fold change in copies of HIV from multivariate model adjusted for age, sex, nadir CD4+ T cell count, timing of ART initiation, ancestry (PCs), and residual variability (probabilistic estimation of expression residuals, PEERs). Bold font denotes genes with q<0.05.
[d] % Change = percent change in host gene expression per two-fold change in copies of HIV

**Supplemental Table 2.** Differentially expressed host genes associated with HIV unspliced RNA (usRNA) in the total study population (top panel) and the European ancestry subgroup (bottom panel), at a Benjamini-Hochberg false discovery rate (FDR) of q<0.25, that were not shown in **Table 2**.

| HIV Unspliced RNA - Total Study Population | | | | | |
|---|---|---|---|---|---|
| **Gene** | **Gene Name** | **p[a]** | **q[b]** | **FC[c]** | **% Change[d]** |
| *MAILR* | Macrophage Interferon Regulatory Lncrna | 4.75E-05 | 0.053 | 0.969 | -3.1 |
| *PLA1A* | Phospholipase A1 Member A | 5.47E-05 | 0.057 | 0.946 | -5.4 |
| *IL6* | Interleukin 6 | 6.41E-05 | 0.062 | 0.926 | -7.4 |
| *PTGES* | Prostaglandin E Synthase | 6.81E-05 | 0.062 | 0.932 | -6.8 |
| *CLEC4D* | C-Type Lectin Domain Family 4 Member D | 6.82E-05 | 0.062 | 0.938 | -6.2 |
| *C15orf48* | Chromosome 15 Open Reading Frame 48 | 7.53E-05 | 0.063 | 0.943 | -5.7 |
| *GBF1* | Golgi Brefeldin A Resistant Guanine Nucleotide Exchange Factor 1 | 7.60E-05 | 0.063 | 1.005 | 0.5 |
| *CLEC4E* | C-Type Lectin Domain Family 4 Member E | 7.88E-05 | 0.063 | 0.927 | -7.3 |
| *AL133163.2* | Novel Transcript | 8.68E-05 | 0.067 | 0.966 | -3.4 |
| *CCRL2* | C-C Motif Chemokine Receptor Like 2 | 9.33E-05 | 0.068 | 0.967 | -3.3 |
| *SLC1A3* | Solute Carrier Family 1 Member 3 | 9.63E-05 | 0.068 | 0.951 | -4.9 |
| *SMPDL3A* | Sphingomyelin Phosphodiesterase Acid Like 3A | 9.90E-05 | 0.068 | 0.937 | -6.3 |
| *CSPG4BP* | Chondroitin Sulfate Proteoglycan Family Member 4B, Pseudogene | 1.00E-04 | 0.072 | 0.952 | -4.8 |
| *AQP9* | Aquaporin 9 | 1.00E-04 | 0.072 | 0.925 | -7.5 |
| *ADM* | Adrenomedullin | 1.00E-04 | 0.075 | 0.950 | -5.0 |
| *TLR8* | Toll Like Receptor 8 | 1.00E-04 | 0.086 | 0.933 | -6.7 |
| *ACOD1* | Aconitate Decarboxylase 1 | 2.00E-04 | 0.086 | 0.941 | -5.9 |
| *LINC02605* | Long Intergenic Non-Protein Coding RNA 2605 | 2.00E-04 | 0.086 | 0.952 | -4.8 |
| *HMGB2* | High Mobility Group Box 2 | 2.00E-04 | 0.086 | 1.007 | 0.7 |
| *FFAR2* | Free Fatty Acid Receptor 2 | 2.00E-04 | 0.086 | 0.932 | -6.8 |
| *SPHK1* | Sphingosine Kinase 1 | 2.00E-04 | 0.086 | 0.966 | -3.4 |
| *RIN2* | Ras And Rab Interactor 2 | 2.00E-04 | 0.089 | 0.938 | -6.2 |
| *F3* | Coagulation Factor III, Tissue Factor | 2.00E-04 | 0.097 | 0.940 | -6.0 |
| *LINC01465* | Long Intergenic Non-Protein Coding RNA 1465 | 2.00E-04 | 0.098 | 0.978 | -2.2 |

| | | | | | |
|---|---|---|---|---|---|
| *PPP1R17* | Protein Phosphatase 1 Regulatory Subunit 17 | 2.00E-04 | 0.101 | 0.954 | -4.6 |
| *OSR2* | Odd-Skipped Related Transciption Factor 2 | 2.00E-04 | 0.101 | 0.950 | -5.0 |
| *AC025580.2* | Novel Transcript, Antisense to SLC30A4 | 2.00E-04 | 0.101 | 0.950 | -5.0 |
| *FPR2* | Formyl Peptide Receptor 2 | 2.00E-04 | 0.101 | 0.936 | -6.4 |
| *TBCC* | Tubulin Folding Cofactor C | 2.00E-04 | 0.102 | 1.010 | 1.0 |
| *VNN3* | Vanin 3 | 2.00E-04 | 0.103 | 0.960 | -4.0 |
| *AL391832.2* | Novel Transcript | 3.00E-04 | 0.111 | 0.958 | -4.2 |
| *CLEC6A* | C-Type Lectin Domain Containing 6A | 3.00E-04 | 0.111 | 0.963 | -3.7 |
| *AC254633.1* | Novel Transcript | 3.00E-04 | 0.111 | 1.037 | 3.7 |
| *C5orf58* | Chromosome 5 Open Reading Frame 58 | 3.00E-04 | 0.111 | 0.984 | -1.6 |
| *DRAM1* | DNA Damage Regulated Autophagy Modulator 1 | 3.00E-04 | 0.111 | 0.970 | -3.0 |
| *SLC22A15* | Solute Carrier Family 22 Member 15 | 3.00E-04 | 0.111 | 0.962 | -3.8 |
| *LINC01093* | Long Intergenic Non-Protein Coding RNA 1093 | 3.00E-04 | 0.111 | 0.954 | -4.6 |
| *ZMAT4* | Zinc Finger Matrin-Type 4 | 3.00E-04 | 0.115 | 0.963 | -3.7 |
| *LSM12* | LSM12 Homolog | 3.00E-04 | 0.115 | 0.994 | -0.6 |
| *VSTM1* | V-Set And Transmembrane Domain Containing 1 | 3.00E-04 | 0.116 | 0.933 | -6.7 |
| *CCL3* | C-C Motif Chemokine Ligand 3 | 3.00E-04 | 0.116 | 0.944 | -5.6 |
| *CLIC4* | Chloride Intracellular Channel 4 | 4.00E-04 | 0.120 | 0.969 | -3.1 |
| *PRLR* | Prolactin Receptor | 4.00E-04 | 0.120 | 0.952 | -4.8 |
| *GPR84* | G Protein-Coupled Receptor 84 | 4.00E-04 | 0.126 | 0.943 | -5.7 |
| *TIGAR* | TP53 Induced Glycolysis Regulatory Phosphatase | 4.00E-04 | 0.126 | 0.988 | -1.2 |
| *SFRP1* | Secreted Frizzled Related Protein 1 | 4.00E-04 | 0.127 | 0.930 | -7.0 |
| *IL1RN* | Interleukin 1 Receptor Antagonist | 4.00E-04 | 0.129 | 0.927 | -7.3 |
| *RAB11FIP3* | RAB11 Family Interacting Protein 3 | 4.00E-04 | 0.130 | 1.006 | 0.6 |
| *PDGFRB* | Platelet Derived Growth Factor Receptor Beta | 4.00E-04 | 0.131 | 0.952 | -4.8 |
| *SLC9A7P1* | Solute Carrier Family 9 Member 7 Pseudogene 1 | 4.00E-04 | 0.131 | 0.946 | -5.4 |
| *TLR4* | Toll Like Receptor 4 | 5.00E-04 | 0.136 | 0.935 | -6.5 |
| *SLC39A1* | Solute Carrier Family 39 Member 1 | 5.00E-04 | 0.141 | 0.973 | -2.7 |
| *CYRIA* | CYFIP Related Rac1 Interactor A | 5.00E-04 | 0.141 | 0.960 | -4.0 |
| *AC037198.1* | Novel Transcript, Sense Intronic to THBS1 | 5.00E-04 | 0.141 | 0.958 | -4.2 |
| *IGSF6* | Immunoglobulin Superfamily Member 6 | 5.00E-04 | 0.142 | 0.981 | -1.9 |
| *ZNF225* | Zinc Finger Protein 225 | 5.00E-04 | 0.142 | 1.013 | 1.3 |

| | | | | | |
|---|---|---|---|---|---|
| *AC245884.11* | Novel Transcript | 5.00E-04 | 0.142 | 0.948 | -5.2 |
| *VAMP2* | Vesicle Associated Membrane Protein 2 | 5.00E-04 | 0.142 | 1.005 | 0.5 |
| *NINJ1* | Ninjurin 1 | 5.00E-04 | 0.142 | 0.982 | -1.8 |
| *CXCL1* | C-X-C Motif Chemokine Ligand 1 | 5.00E-04 | 0.142 | 0.943 | -5.7 |
| *KTI12* | KTI12 Chromatin Associated Homolog | 6.00E-04 | 0.144 | 1.009 | 0.9 |
| *FXYD6* | FXYD Domain Containing Ion Transport Regulator 6 | 6.00E-04 | 0.145 | 0.952 | -4.8 |
| *UICLM* | Up-Regulated In Colorectal Cancer Liver Metastasis | 6.00E-04 | 0.151 | 0.948 | -5.2 |
| *ARRDC3* | Arrestin Domain Containing 3 | 6.00E-04 | 0.154 | 1.013 | 1.3 |
| *FRMD7* | FERM Domain Containing 7 | 6.00E-04 | 0.154 | 0.955 | -4.5 |
| *MEFV* | MEFV Innate Immuity Regulator, Pyrin | 7.00E-04 | 0.156 | 0.946 | -5.4 |
| *SPIB* | Spi-B Transcription Factor | 7.00E-04 | 0.156 | 0.954 | -4.6 |
| *HCAR3* | Hydroxycarboxylic Acid Receptor 3 | 7.00E-04 | 0.167 | 0.944 | -5.6 |
| *MTCYBP23* | MT-CYB Pseudogene 23 | 7.00E-04 | 0.168 | 0.975 | -2.5 |
| *C1QA* | Complement C1q A Chain | 7.00E-04 | 0.169 | 0.944 | -5.6 |
| *VCAM1* | Vascular Cell Adhesion Molecule 1 | 8.00E-04 | 0.172 | 0.952 | -4.8 |
| *AL117335.1* | Novel Transcript, Antisense to SIRPA | 8.00E-04 | 0.181 | 0.966 | -3.4 |
| *BNC2* | Basonuclin 2 | 8.00E-04 | 0.181 | 0.952 | -4.8 |
| *IRAK2* | Interleukin 1 Receptor Associated Kinase 2 | 8.00E-04 | 0.181 | 0.977 | -2.3 |
| *AC112128.1* | Novel Protein | 8.00E-04 | 0.183 | 1.016 | 1.6 |
| *RPH3A* | Rabphilin 3A | 9.00E-04 | 0.189 | 0.950 | -5.0 |
| *AC011990.1* | Novel Transcript | 9.00E-04 | 0.194 | 0.968 | -3.2 |
| *NFKBIZ* | NFKB Inhibitor Zeta | 9.00E-04 | 0.195 | 0.983 | -1.7 |
| *LINC02413* | Long Intergenic Non-Protein Coding RNA 2413 ( | 9.00E-04 | 0.196 | 0.956 | -4.4 |
| *MYO6* | Myosin VI | 1.00E-03 | 0.196 | 0.982 | -1.8 |
| *AL353593.1* | Novel Transcript, Antisense to OBSCN | 1.00E-03 | 0.198 | 0.987 | -1.3 |
| *AC079753.2* | Novel Transcript | 1.00E-03 | 0.202 | 0.957 | -4.3 |
| *AC106881.1* | Novel Transcript, Antisense to UNC5C | 1.00E-03 | 0.202 | 0.967 | -3.3 |
| *RNF144B* | Ring Finger Protein 144B | 1.00E-03 | 0.202 | 0.970 | -3.0 |
| *ADPRH* | ADP-Ribosylarginine Hydrolase | 1.00E-03 | 0.202 | 0.981 | -1.9 |
| *IFNB1* | Interferon Beta 1 | 1.00E-03 | 0.203 | 0.950 | -5.0 |
| *LPL* | Lipoprotein Lipase | 1.10E-03 | 0.205 | 0.956 | -4.4 |
| *RPS15AP30* | Ribosomal Protein S15a Pseudogene 30 | 1.10E-03 | 0.217 | 0.961 | -3.9 |
| *SCARF1* | Scavenger Receptor Class F Member 1 | 1.20E-03 | 0.222 | 0.980 | -2.0 |

| | | | | | |
|---|---|---|---|---|---|
| *C4orf46* | Chromosome 4 Open Reading Frame 46 | 1.20E-03 | 0.222 | 0.989 | -1.1 |
| *AC096733.2* | Novel Transcript | 1.20E-03 | 0.222 | 0.953 | -4.7 |
| *AC115618.1* | Novel Transcript, Antisense to RBM3 | 1.20E-03 | 0.226 | 0.974 | -2.6 |
| *TCF7L2* | Transcription Factor 7 Like 2 | 1.30E-03 | 0.228 | 0.984 | -1.6 |
| *CCL3L3* | C-C Motif Chemokine Ligand 3 Like 3 | 1.30E-03 | 0.234 | 0.921 | -7.9 |
| *CCL20* | C-C Motif Chemokine Ligand 20 | 1.40E-03 | 0.246 | 0.965 | -3.5 |
| *MAFF* | MAF Bzip Transcription Factor F | 1.40E-03 | 0.246 | 0.984 | -1.6 |
| *KRT5* | Keratin 5 | 1.40E-03 | 0.248 | 0.954 | -4.6 |
| **HIV Unspliced RNA – European Ancestry Subgroup** | | | | | |
| **Gene** | **Gene Name** | **p[a]** | **q[b]** | **FC[c]** | **% Change[d]** |
| *MTCYBP23* | MT-CYB Pseudogene 23 | 1.88E-05 | 0.052 | 0.959 | -4.1 |
| *ACOD1* | Aconitate Decarboxylase 1 | 2.35E-05 | 0.057 | 0.919 | -8.1 |
| *ZMAT4* | Zinc Finger Matrin-Type 4 | 2.81E-05 | 0.060 | 0.941 | -5.9 |
| *IFNB1* | Interferon Beta 1 | 3.35E-05 | 0.065 | 0.921 | -7.9 |
| *MAILR* | Macrophage Interferon Regulatory Lncrna | 4.49E-05 | 0.079 | 0.958 | -4.2 |
| *GHITM* | Growth Hormone Inducible Transmembrane Protein | 6.25E-05 | 0.096 | 0.993 | -0.7 |
| *NBN* | Nibrin | 7.23E-05 | 0.096 | 0.989 | -1.1 |
| *SFRP1* | Secreted Frizzled Related Protein 1 | 7.46E-05 | 0.096 | 0.881 | -11.9 |
| *KCNJ2-AS1* | KCNJ2 Antisense RNA 1 | 7.48E-05 | 0.096 | 0.900 | -10.0 |
| *DAPK1-IT1* | DAPK1 Intronic Transcript 1 | 8.18E-05 | 0.099 | 0.933 | -6.7 |
| *MMP2-AS1* | MMP2 Antisense RNA 1 | 1.00E-04 | 0.116 | 0.937 | -6.3 |
| *AC106712.1* | Novel Transcript | 1.00E-04 | 0.116 | 0.950 | -5.0 |
| *LINC00211* | Long Intergenic Non-Protein Coding RNA 211 | 1.00E-04 | 0.116 | 0.946 | -5.4 |
| *CCRL2* | C-C Motif Chemokine Receptor Like 2 | 1.00E-04 | 0.121 | 0.960 | -4.0 |
| *RPL36AP16* | Ribosomal Protein L36a Pseudogene 16 | 2.00E-04 | 0.152 | 1.030 | 3.0 |
| *CSPG4BP* | Chondroitin Sulfate Proteoglycan Family Member 4B, Pseudogene | 2.00E-04 | 0.164 | 0.934 | -6.6 |
| *AC091173.1* | Novel Transcript | 2.00E-04 | 0.173 | 0.952 | -4.8 |
| *SLC39A1* | Solute Carrier Family 39 Member 1 | 2.00E-04 | 0.179 | 0.960 | -4.0 |
| *TCF7L2* | Transcription Factor 7 Like 2 | 2.00E-04 | 0.181 | 0.979 | -2.1 |
| *ADPRH* | ADP-Ribosylarginine Hydrolase | 3.00E-04 | 0.189 | 0.973 | -2.7 |
| *MIR3945HG* | MIR3945 Host Gene | 3.00E-04 | 0.189 | 0.935 | -6.5 |
| *CSF3* | Colony Stimulating Factor 3 | 3.00E-04 | 0.189 | 0.913 | -8.7 |
| *BNC2* | Basonuclin 2 | 3.00E-04 | 0.189 | 0.932 | -6.8 |
| *AC096733.2* | Novel Transcript | 3.00E-04 | 0.189 | 0.929 | -7.1 |

| | | | | | |
|---|---|---|---|---|---|
| *IL1A* | Interleukin 1 Alpha | 3.00E-04 | 0.191 | 0.898 | -10.2 |
| *AC106881.1* | Novel Transcript, Antisense to UNC5C | 4.00E-04 | 0.215 | 0.954 | -4.6 |
| *KTI12* | KTI12 Chromatin Associated Homolog | 4.00E-04 | 0.215 | 1.013 | 1.3 |
| *ZNF366* | Zinc Finger Protein 366 | 4.00E-04 | 0.215 | 1.070 | 7.0 |
| *LINC00677* | Long Intergenic Non-Protein Coding RNA 677 | 4.00E-04 | 0.215 | 0.952 | -4.8 |
| *LPL* | Lipoprotein Lipase | 4.00E-04 | 0.218 | 0.940 | -6.0 |
| *AC092723.1* | Novel Transcript | 4.00E-04 | 0.232 | 0.961 | -3.9 |
| *AC011990.1* | Novel Transcript | 5.00E-04 | 0.232 | 0.955 | -4.5 |
| *SLC22A15* | Solute Carrier Family 22 Member 15 | 5.00E-04 | 0.234 | 0.951 | -4.9 |
| *AC096667.1* | Proline-Rich Protein 18-Like | 5.00E-04 | 0.240 | 0.936 | -6.4 |
| *RRN3P4* | RRN3 Pseudogene 4 | 5.00E-04 | 0.240 | 0.942 | -5.8 |
| *RNU1-103P* | RNA, U1 Small Nuclear 103, Pseudogene | 5.00E-04 | 0.240 | 0.976 | -2.4 |
| *CXCL3* | C-X-C Motif Chemokine Ligand 3 | 5.00E-04 | 0.240 | 0.915 | -8.5 |
| *IMP3* | IMP U3 Small Nucleolar Ribonucleoprotein 3 | 6.00E-04 | 0.248 | 1.013 | 1.3 |
| *KCNJ2* | Potassium Inwardly Rectifying Channel Subfamily J Member 2 | 6.00E-04 | 0.248 | 0.906 | -9.4 |
| *OTUD3* | OTU Deubiquitinase 3 | 6.00E-04 | 0.248 | 0.990 | -1.0 |
| *VNN3* | Vanin 3 (Source:HGNC Symbol;Acc:HGNC:16431) | 7.00E-04 | 0.248 | 0.955 | -4.5 |
| *AC066616.2* | Mitochondrially Encoded NADH 4L (MT-ND4L) Pseudogene | 7.00E-04 | 0.248 | 0.967 | -3.3 |
| *THBD* | Thrombomodulin | 7.00E-04 | 0.248 | 1.063 | 6.3 |
| *ZNF563* | Zinc Finger Protein 563 | 7.00E-04 | 0.248 | 1.016 | 1.6 |
| *PRLR* | Prolactin Receptor | 7.00E-04 | 0.248 | 0.938 | -6.2 |
| *TECPR2* | Tectonin Beta-Propeller Repeat Containing 2 | 7.00E-04 | 0.248 | 0.990 | -1.0 |
| *TUSC3* | Tumor Suppressor Candidate 3 | 7.00E-04 | 0.248 | 1.094 | 9.4 |

[a] p = two sided p-value.
[b] q = two-sided false discovery rate (FDR) Benjamini-Hochberg q-value.
[c] FC = fold-change in host gene expression per two-fold change in copies of HIV from multivariate model adjusted for age, sex, nadir CD4+ T cell count, timing of ART initiation, ancestry (PCs), and residual variability (probabilistic estimation of expression residuals, PEERs).
[d] % Change = percent change in host gene expression per two-fold change in copies of HIV.

**Supplemental Table 3.** Gene set enrichment analyses (GSEA) of ranked differentially expressed genes in relation to HIV total DNA using the Gene Ontology Biological Processes (GO-BP) database. Genes sets with Benjamini-Hochberg false discovery rate (FDR)-adjusted q<0.25 are shown for the total study population (top panel) and for the European ancestry subgroup (bottom panel). Gene sets where q<0.05 are shown in bold font.

| HIV Total DNA | | | | | |
|---|---|---|---|---|---|
| | **GO ID** | **Description** | **NES[a]** | **p[b]** | **q[c]** |
| **Total Study Population** | | | | | |
| 1 | GO:0036507 | protein demannosylation | 1.9 | 6.04E-05 | 0.171 |
| 2 | GO:0036508 | protein alpha-1,2-demannosylation | 1.9 | 6.04E-05 | 0.171 |
| 3 | GO:0006333 | chromatin assembly or disassembly | 1.4 | 1.00E-04 | 0.211 |
| **European Ancestry Subgroup** | | | | | |
| 1 | GO:0006958 | complement activation, classical pathway | 1.8 | 2.26E-10 | **1.27E-06** |
| 2 | GO:0030449 | regulation of complement activation | 1.9 | 3.50E-09 | **9.80E-06** |
| 3 | GO:0006956 | complement activation | 1.7 | 6.65E-09 | **1.24E-05** |
| 4 | GO:0002920 | regulation of humoral immune response | 1.8 | 8.98E-09 | **1.26E-05** |
| 5 | GO:0002455 | humoral immune response mediated by circulating immunoglobulin | 1.7 | 1.44E-08 | **1.61E-05** |
| 6 | GO:0006959 | humoral immune response | 1.5 | 9.50E-07 | **9.00E-04** |
| 7 | GO:0019724 | B cell mediated immunity | 1.5 | 2.36E-06 | **0.002** |
| 8 | GO:0016064 | immunoglobulin mediated immune response | 1.5 | 3.91E-06 | **0.003** |
| 9 | GO:0002449 | lymphocyte mediated immunity | 1.4 | 4.74E-06 | **0.003** |
| 10 | GO:0006910 | phagocytosis, recognition | 1.7 | 6.19E-06 | **0.004** |
| 11 | GO:0006911 | phagocytosis, engulfment | 1.6 | 1.04E-05 | **0.005** |
| 12 | GO:0002460 | adaptive immune response based on somatic recombination of immune receptors built from immunoglobulin superfamily domains | 1.4 | 1.66E-05 | **0.008** |
| 13 | GO:0099024 | plasma membrane invagination | 1.6 | 4.18E-05 | **0.018** |
| 14 | GO:0038094 | Fc-gamma receptor signaling pathway | 1.5 | 5.58E-05 | **0.020** |
| 15 | GO:0002433 | immune response-regulating cell surface receptor signaling pathway involved in phagocytosis | 1.5 | 5.64E-05 | **0.020** |
| 16 | GO:0038096 | Fc-gamma receptor signaling pathway involved in phagocytosis | 1.5 | 5.64E-05 | **0.020** |
| 17 | GO:0042742 | defense response to bacterium | 1.4 | 8.86E-05 | **0.029** |
| 18 | GO:0010324 | membrane invagination | 1.5 | 1.00E-04 | **0.034** |
| 19 | GO:0002431 | Fc receptor mediated stimulatory signaling pathway | 1.5 | 1.00E-04 | **0.040** |
| 20 | GO:0002377 | immunoglobulin production | 1.4 | 1.00E-04 | **0.041** |
| 21 | GO:0070268 | cornification | 1.7 | 2.00E-04 | 0.066 |
| 22 | GO:0043032 | positive regulation of macrophage activation | 1.8 | 6.00E-04 | 0.141 |
| 23 | GO:0038095 | Fc-epsilon receptor signaling pathway | 1.4 | 9.00E-04 | 0.231 |

| | | | | | |
|---|---|---|---|---|---|
| 24 | GO:0002440 | production of molecular mediator of immune response | 1.3 | 1.00E-03 | 0.238 |
| 25 | GO:0002697 | regulation of immune effector process | 1.2 | 1.10E-03 | 0.245 |
| 26 | GO:0010463 | mesenchymal cell proliferation | 1.7 | 1.20E-03 | 0.248 |

[a] NES = normalized enrichment score.
[b] p = two sided p-value.
[c] q = two-sided false discovery rate (FDR) Benjamini-Hochberg q-value.

**Supplemental Table 4.** Gene set enrichment analyses (GSEA) of ranked differentially expressed genes in relation to HIV unspliced RNA using the Gene Ontology Biological Processes (GO-BP) database. Genes sets with Benjamini-Hochberg false discovery rate (FDR)-adjusted q<0.05 are shown for the total study population (top panel) and for the European ancestry subgroup (bottom panel). Gene sets where q<0.05 are shown in bold font.

| **HIV Unspliced RNA - Total Study Population** | | | | | |
|---|---|---|---|---|---|
| **Rank** | **GO ID** | **Description** | **NES[a]** | **p[b]** | **q[c]** |
| 1 | GO:0009617 | response to bacterium | 1.4 | 1.34E-08 | **7.55E-05** |
| 2 | GO:0031663 | lipopolysaccharide-mediated signaling pathway | 1.8 | 2.05E-06 | **0.006** |
| 3 | GO:0071222 | cellular response to lipopolysaccharide | 1.5 | 3.35E-06 | **0.006** |
| 4 | GO:0032640 | tumor necrosis factor production | 1.6 | 4.39E-06 | **0.006** |
| 5 | GO:0001818 | negative regulation of cytokine production | 1.4 | 4.87E-06 | **0.006** |
| 6 | GO:0071706 | tumor necrosis factor superfamily cytokine production | 1.5 | 6.46E-06 | **0.006** |
| 7 | GO:0032611 | interleukin-1 beta production | 1.6 | 9.80E-06 | **0.008** |
| 8 | GO:0071219 | cellular response to molecule of bacterial origin | 1.5 | 1.34E-05 | **0.008** |
| 9 | GO:0032652 | regulation of interleukin-1 production | 1.6 | 1.55E-05 | **0.008** |
| 10 | GO:0032680 | regulation of tumor necrosis factor production | 1.5 | 1.61E-05 | **0.008** |
| 11 | GO:0032651 | regulation of interleukin-1 beta production | 1.7 | 1.75E-05 | **0.008** |
| 12 | GO:0071216 | cellular response to biotic stimulus | 1.4 | 1.75E-05 | **0.008** |
| 13 | GO:1903555 | regulation of tumor necrosis factor superfamily cytokine production | 1.5 | 1.75E-05 | **0.008** |
| 14 | GO:0002237 | response to molecule of bacterial origin | 1.4 | 2.08E-05 | **0.008** |
| 15 | GO:0032496 | response to lipopolysaccharide | 1.4 | 2.42E-05 | **0.009** |
| 16 | GO:0002697 | regulation of immune effector process | 1.3 | 3.17E-05 | **0.011** |
| 17 | GO:0042108 | positive regulation of cytokine biosynthetic process | 1.7 | 4.35E-05 | **0.014** |
| 18 | GO:0036230 | granulocyte activation | 1.3 | 4.52E-05 | **0.014** |
| 19 | GO:0033002 | muscle cell proliferation | 1.5 | 5.81E-05 | **0.017** |
| 20 | GO:0032612 | interleukin-1 production | 1.6 | 6.26E-05 | **0.018** |
| 21 | GO:0032635 | interleukin-6 production | 1.5 | 7.16E-05 | **0.019** |
| 22 | GO:0071396 | cellular response to lipid | 1.3 | 7.84E-05 | **0.020** |
| 23 | GO:1904646 | cellular response to amyloid-beta | 1.8 | 8.29E-05 | **0.020** |
| 24 | GO:0002275 | myeloid cell activation involved in immune response | 1.3 | 8.40E-05 | **0.020** |
| 25 | GO:0006959 | humoral immune response | 1.4 | 9.76E-05 | **0.022** |
| 26 | GO:0042119 | neutrophil activation | 1.3 | 1.00E-04 | **0.022** |
| 27 | GO:0032649 | regulation of interferon-gamma production | 1.6 | 1.00E-04 | **0.022** |
| 28 | GO:1904645 | response to amyloid-beta | 1.8 | 1.00E-04 | **0.022** |
| 29 | GO:0002827 | positive regulation of T-helper 1 type immune response | 1.9 | 1.00E-04 | **0.022** |
| 30 | GO:1904037 | positive regulation of epithelial cell apoptotic process | 1.7 | 1.00E-04 | **0.022** |
| 31 | GO:0043312 | neutrophil degranulation | 1.3 | 1.00E-04 | **0.022** |

| | | | | | |
|---|---|---|---|---|---|
| 32 | GO:0050688 | regulation of defense response to virus | 1.6 | 1.00E-04 | **0.023** |
| 33 | GO:0060428 | lung epithelium development | 1.9 | 1.00E-04 | **0.023** |
| 34 | GO:0002444 | myeloid leukocyte mediated immunity | 1.3 | 1.00E-04 | **0.023** |
| 35 | GO:0002221 | pattern recognition receptor signaling pathway | 1.4 | 2.00E-04 | **0.024** |
| 36 | GO:0050691 | regulation of defense response to virus by host | 1.7 | 2.00E-04 | **0.024** |
| 37 | GO:0002446 | neutrophil mediated immunity | 1.3 | 2.00E-04 | **0.024** |
| 38 | GO:0007249 | I-kappaB kinase/NF-kappaB signaling | 1.4 | 2.00E-04 | **0.024** |
| 39 | GO:0038094 | Fc-gamma receptor signaling pathway | 1.4 | 2.00E-04 | **0.024** |
| 40 | GO:0060740 | prostate gland epithelium morphogenesis | 2.0 | 2.00E-04 | **0.025** |
| 41 | GO:0032755 | positive regulation of interleukin-6 production | 1.6 | 2.00E-04 | **0.025** |
| 42 | GO:0002283 | neutrophil activation involved in immune response | 1.3 | 2.00E-04 | **0.025** |
| 43 | GO:0002224 | toll-like receptor signaling pathway | 1.4 | 2.00E-04 | **0.025** |
| 44 | GO:0042035 | regulation of cytokine biosynthetic process | 1.5 | 2.00E-04 | **0.026** |
| 45 | GO:0032675 | regulation of interleukin-6 production | 1.5 | 2.00E-04 | **0.031** |
| 46 | GO:0043299 | leukocyte degranulation | 1.3 | 2.00E-04 | **0.031** |
| 47 | GO:0071674 | mononuclear cell migration | 1.6 | 3.00E-04 | **0.032** |
| 48 | GO:0008544 | epidermis development | 1.4 | 3.00E-04 | **0.033** |
| 49 | GO:0006909 | phagocytosis | 1.3 | 3.00E-04 | **0.034** |
| 50 | GO:0032731 | positive regulation of interleukin-1 beta production | 1.6 | 3.00E-04 | **0.037** |
| 51 | GO:0032653 | regulation of interleukin-10 production | 1.6 | 3.00E-04 | **0.037** |
| 52 | GO:0043331 | response to dsRNA | 1.7 | 3.00E-04 | **0.038** |
| 53 | GO:0001773 | myeloid dendritic cell activation | 1.8 | 4.00E-04 | **0.040** |
| 54 | GO:0009913 | epidermal cell differentiation | 1.4 | 4.00E-04 | **0.041** |
| 55 | GO:0032613 | interleukin-10 production | 1.6 | 4.00E-04 | **0.041** |
| 56 | GO:0060512 | prostate gland morphogenesis | 1.8 | 4.00E-04 | **0.042** |
| 57 | GO:0097028 | dendritic cell differentiation | 1.7 | 4.00E-04 | **0.042** |
| 58 | GO:0030216 | keratinocyte differentiation | 1.4 | 4.00E-04 | **0.042** |
| 59 | GO:0006898 | receptor-mediated endocytosis | 1.3 | 5.00E-04 | **0.043** |
| 60 | GO:0032732 | positive regulation of interleukin-1 production | 1.6 | 5.00E-04 | **0.043** |
| 61 | GO:0150077 | regulation of neuroinflammatory response | 1.8 | 5.00E-04 | **0.043** |
| 62 | GO:0050663 | cytokine secretion | 1.4 | 5.00E-04 | **0.043** |
| 63 | GO:0050900 | leukocyte migration | 1.2 | 5.00E-04 | **0.043** |
| 64 | GO:0002431 | Fc receptor mediated stimulatory signaling pathway | 1.4 | 5.00E-04 | **0.043** |
| 65 | GO:1903557 | positive regulation of tumor necrosis factor superfamily cytokine production | 1.5 | 5.00E-04 | **0.043** |
| 66 | GO:0002460 | adaptive immune response based on somatic recombination of immune receptors built from immunoglobulin superfamily domains | 1.3 | 5.00E-04 | **0.044** |
| 67 | GO:0002548 | monocyte chemotaxis | 1.6 | 5.00E-04 | **0.044** |
| 68 | GO:0043588 | skin development | 1.4 | 5.00E-04 | **0.044** |
| 69 | GO:0032760 | positive regulation of tumor necrosis factor production | 1.5 | 6.00E-04 | **0.046** |
| 70 | GO:0050727 | regulation of inflammatory response | 1.3 | 6.00E-04 | **0.046** |

| | | | | | |
|---|---|---|---|---|---|
| 71 | GO:0006816 | calcium ion transport | 1.3 | 6.00E-04 | **0.046** |
| 72 | GO:0002433 | immune response-regulating cell surface receptor signaling pathway involved in phagocytosis | 1.4 | 6.00E-04 | **0.046** |
| 73 | GO:0038096 | Fc-gamma receptor signaling pathway involved in phagocytosis | 1.4 | 6.00E-04 | **0.046** |
| 74 | GO:0030595 | leukocyte chemotaxis | 1.4 | 6.00E-04 | **0.047** |
| 75 | GO:0032602 | chemokine production | 1.5 | 6.00E-04 | **0.049** |
| 76 | GO:0002819 | regulation of adaptive immune response | 1.4 | 7.00E-04 | **0.049** |

### HIV Unspliced RNA - European Ancestry Subgroup

| Rank | GO ID | Description | NES[a] | p[b] | q[c] |
|---|---|---|---|---|---|
| 1 | GO:0009617 | response to bacterium | 1.5 | 3.55E-12 | **1.99E-08** |
| 2 | GO:0001819 | positive regulation of cytokine production | 1.5 | 1.97E-09 | **5.50E-06** |
| 3 | GO:0032496 | response to lipopolysaccharide | 1.5 | 7.04E-09 | **1.31E-05** |
| 4 | GO:0002237 | response to molecule of bacterial origin | 1.5 | 9.88E-09 | **1.38E-05** |
| 5 | GO:0032103 | positive regulation of response to external stimulus | 1.4 | 2.48E-08 | **2.78E-05** |
| 6 | GO:0031349 | positive regulation of defense response | 1.5 | 4.93E-08 | **4.60E-05** |
| 7 | GO:0002699 | positive regulation of immune effector process | 1.5 | 3.24E-07 | **3.00E-04** |
| 8 | GO:0023061 | signal release | 1.3 | 4.47E-07 | **3.00E-04** |
| 9 | GO:0002694 | regulation of leukocyte activation | 1.3 | 1.24E-06 | **8.00E-04** |
| 10 | GO:0050729 | positive regulation of inflammatory response | 1.6 | 1.43E-06 | **8.00E-04** |
| 11 | GO:0071216 | cellular response to biotic stimulus | 1.5 | 1.72E-06 | **9.00E-04** |
| 12 | GO:0050727 | regulation of inflammatory response | 1.4 | 1.91E-06 | **9.00E-04** |
| 13 | GO:0002449 | lymphocyte mediated immunity | 1.4 | 2.94E-06 | **0.001** |
| 14 | GO:0071222 | cellular response to lipopolysaccharide | 1.5 | 4.94E-06 | **0.002** |
| 15 | GO:0022407 | regulation of cell-cell adhesion | 1.4 | 5.29E-06 | **0.002** |
| 16 | GO:0071219 | cellular response to molecule of bacterial origin | 1.5 | 6.12E-06 | **0.002** |
| 17 | GO:0002703 | regulation of leukocyte mediated immunity | 1.5 | 6.74E-06 | **0.002** |
| 18 | GO:0002708 | positive regulation of lymphocyte mediated immunity | 1.6 | 7.29E-06 | **0.002** |
| 19 | GO:0035747 | natural killer cell chemotaxis | 2.1 | 7.29E-06 | **0.002** |
| 20 | GO:0050663 | cytokine secretion | 1.5 | 1.24E-05 | **0.003** |
| 21 | GO:0035743 | CD4-positive, alpha-beta T cell cytokine production | 2.0 | 1.29E-05 | **0.003** |
| 22 | GO:0050715 | positive regulation of cytokine secretion | 1.6 | 1.29E-05 | **0.003** |
| 23 | GO:0002292 | T cell differentiation involved in immune response | 1.7 | 1.30E-05 | **0.003** |
| 24 | GO:0045622 | regulation of T-helper cell differentiation | 1.9 | 1.31E-05 | **0.003** |
| 25 | GO:0002367 | cytokine production involved in immune response | 1.6 | 1.34E-05 | **0.003** |
| 26 | GO:0031663 | lipopolysaccharide-mediated signaling pathway | 1.7 | 1.61E-05 | **0.004** |
| 27 | GO:0002705 | positive regulation of leukocyte mediated immunity | 1.5 | 1.77E-05 | **0.004** |
| 28 | GO:1904646 | cellular response to amyloid-beta | 1.8 | 1.97E-05 | **0.004** |
| 29 | GO:0002548 | monocyte chemotaxis | 1.8 | 2.00E-05 | **0.004** |
| 30 | GO:0002706 | regulation of lymphocyte mediated immunity | 1.5 | 2.11E-05 | **0.004** |
| 31 | GO:0002460 | adaptive immune response based on somatic recombination of immune receptors built from immunoglobulin superfamily domains | 1.3 | 2.16E-05 | **0.004** |

| | | | | | |
|---|---|---|---|---|---|
| 32 | GO:0043370 | regulation of CD4-positive, alpha-beta T cell differentiation | 1.8 | 2.22E-05 | **0.004** |
| 33 | GO:0055057 | neuroblast division | 2.1 | 2.61E-05 | **0.004** |
| 34 | GO:0050905 | neuromuscular process | 1.7 | 2.69E-05 | **0.004** |
| 35 | GO:0042093 | T-helper cell differentiation | 1.7 | 2.75E-05 | **0.004** |
| 36 | GO:0050707 | regulation of cytokine secretion | 1.5 | 2.75E-05 | **0.004** |
| 37 | GO:0043367 | CD4-positive, alpha-beta T cell differentiation | 1.6 | 2.89E-05 | **0.004** |
| 38 | GO:0032649 | regulation of interferon-gamma production | 1.6 | 3.06E-05 | **0.005** |
| 39 | GO:0070374 | positive regulation of ERK1 and ERK2 cascade | 1.5 | 3.17E-05 | **0.005** |
| 40 | GO:0043410 | positive regulation of MAPK cascade | 1.3 | 3.96E-05 | **0.006** |
| 41 | GO:0002718 | regulation of cytokine production involved in immune response | 1.6 | 4.29E-05 | **0.006** |
| 42 | GO:0032755 | positive regulation of interleukin-6 production | 1.6 | 4.35E-05 | **0.006** |
| 43 | GO:0050863 | regulation of T cell activation | 1.4 | 4.35E-05 | **0.006** |
| 44 | GO:2000514 | regulation of CD4-positive, alpha-beta T cell activation | 1.7 | 4.46E-05 | **0.006** |
| 45 | GO:0002286 | T cell activation involved in immune response | 1.6 | 4.74E-05 | **0.006** |
| 46 | GO:0030593 | neutrophil chemotaxis | 1.6 | 4.74E-05 | **0.006** |
| 47 | GO:0060191 | regulation of lipase activity | 1.6 | 4.80E-05 | **0.006** |
| 48 | GO:0051249 | regulation of lymphocyte activation | 1.3 | 4.91E-05 | **0.006** |
| 49 | GO:0061900 | glial cell activation | 1.7 | 4.91E-05 | **0.006** |
| 50 | GO:0097529 | myeloid leukocyte migration | 1.4 | 4.91E-05 | **0.006** |
| 51 | GO:0002720 | positive regulation of cytokine production involved in immune response | 1.7 | 5.53E-05 | **0.006** |
| 52 | GO:0032635 | interleukin-6 production | 1.5 | 5.64E-05 | **0.006** |
| 53 | GO:0060193 | positive regulation of lipase activity | 1.7 | 5.64E-05 | **0.006** |
| 54 | GO:0060326 | cell chemotaxis | 1.4 | 5.70E-05 | **0.006** |
| 55 | GO:0001906 | cell killing | 1.5 | 5.81E-05 | **0.006** |
| 56 | GO:0042742 | defense response to bacterium | 1.4 | 6.26E-05 | **0.006** |
| 57 | GO:0003229 | ventricular cardiac muscle tissue development | 1.8 | 6.37E-05 | **0.006** |
| 58 | GO:0032609 | interferon-gamma production | 1.5 | 6.71E-05 | **0.006** |
| 59 | GO:0051770 | positive regulation of nitric-oxide synthase biosynthetic process | 2.0 | 6.82E-05 | **0.006** |
| 60 | GO:0042100 | B cell proliferation | 1.6 | 6.94E-05 | **0.006** |
| 61 | GO:1903037 | regulation of leukocyte cell-cell adhesion | 1.4 | 6.94E-05 | **0.006** |
| 62 | GO:0002526 | acute inflammatory response | 1.6 | 7.05E-05 | **0.006** |
| 63 | GO:0007159 | leukocyte cell-cell adhesion | 1.3 | 7.05E-05 | **0.006** |
| 64 | GO:0050867 | positive regulation of cell activation | 1.3 | 7.28E-05 | **0.006** |
| 65 | GO:0032611 | interleukin-1 beta production | 1.6 | 7.39E-05 | **0.006** |
| 66 | GO:0033002 | muscle cell proliferation | 1.4 | 7.50E-05 | **0.006** |
| 67 | GO:0002697 | regulation of immune effector process | 1.3 | 7.61E-05 | **0.006** |
| 68 | GO:0090179 | planar cell polarity pathway involved in neural tube closure | 2.0 | 7.61E-05 | **0.006** |
| 69 | GO:0001505 | regulation of neurotransmitter levels | 1.4 | 7.95E-05 | **0.006** |
| 70 | GO:1902106 | negative regulation of leukocyte differentiation | 1.6 | 7.95E-05 | **0.006** |

| | | | | | |
|---|---|---|---|---|---|
| 71 | GO:0002369 | T cell cytokine production | 1.7 | 8.29E-05 | **0.007** |
| 72 | GO:0042108 | positive regulation of cytokine biosynthetic process | 1.6 | 8.40E-05 | **0.007** |
| 73 | GO:0032675 | regulation of interleukin-6 production | 1.5 | 9.19E-05 | **0.007** |
| 74 | GO:0042116 | macrophage activation | 1.6 | 9.19E-05 | **0.007** |
| 75 | GO:0002695 | negative regulation of leukocyte activation | 1.4 | 9.53E-05 | **0.007** |
| 76 | GO:0001909 | leukocyte mediated cytotoxicity | 1.5 | 9.64E-05 | **0.007** |
| 77 | GO:0070372 | regulation of ERK1 and ERK2 cascade | 1.4 | 9.76E-05 | **0.007** |
| 78 | GO:0007249 | I-kappaB kinase/NF-kappaB signaling | 1.3 | 1.00E-04 | **0.008** |
| 79 | GO:0001910 | regulation of leukocyte mediated cytotoxicity | 1.6 | 1.00E-04 | **0.008** |
| 80 | GO:0001774 | microglial cell activation | 1.7 | 1.00E-04 | **0.008** |
| 81 | GO:0002269 | leukocyte activation involved in inflammatory response | 1.7 | 1.00E-04 | **0.008** |
| 82 | GO:0150076 | neuroinflammatory response | 1.6 | 1.00E-04 | **0.008** |
| 83 | GO:0030595 | leukocyte chemotaxis | 1.4 | 1.00E-04 | **0.009** |
| 84 | GO:0070371 | ERK1 and ERK2 cascade | 1.4 | 1.00E-04 | **0.009** |
| 85 | GO:0002294 | CD4-positive, alpha-beta T cell differentiation involved in immune response | 1.6 | 1.00E-04 | **0.009** |
| 86 | GO:0009615 | response to virus | 1.3 | 1.00E-04 | **0.009** |
| 87 | GO:0031341 | regulation of cell killing | 1.5 | 1.00E-04 | **0.009** |
| 88 | GO:0002456 | T cell mediated immunity | 1.5 | 1.00E-04 | **0.009** |
| 89 | GO:0045620 | negative regulation of lymphocyte differentiation | 1.7 | 2.00E-04 | **0.010** |
| 90 | GO:0032612 | interleukin-1 production | 1.5 | 2.00E-04 | **0.010** |
| 91 | GO:0000186 | activation of MAPKK activity | 1.7 | 2.00E-04 | **0.010** |
| 92 | GO:1905276 | regulation of epithelial tube formation | 1.9 | 2.00E-04 | **0.011** |
| 93 | GO:0042330 | taxis | 1.2 | 2.00E-04 | **0.011** |
| 94 | GO:0050708 | regulation of protein secretion | 1.3 | 2.00E-04 | **0.011** |
| 95 | GO:0045089 | positive regulation of innate immune response | 1.4 | 2.00E-04 | **0.012** |
| 96 | GO:0071396 | cellular response to lipid | 1.2 | 2.00E-04 | **0.012** |
| 97 | GO:0097696 | receptor signaling pathway via STAT | 1.5 | 2.00E-04 | **0.012** |
| 98 | GO:1902105 | regulation of leukocyte differentiation | 1.3 | 2.00E-04 | **0.012** |
| 99 | GO:0031295 | T cell costimulation | 1.6 | 2.00E-04 | **0.013** |
| 100 | GO:0035710 | CD4-positive, alpha-beta T cell activation | 1.5 | 2.00E-04 | **0.013** |
| 101 | GO:0045581 | negative regulation of T cell differentiation | 1.7 | 3.00E-04 | **0.014** |
| 102 | GO:0042089 | cytokine biosynthetic process | 1.5 | 3.00E-04 | **0.015** |
| 103 | GO:0046688 | response to copper ion | 1.8 | 3.00E-04 | **0.015** |
| 104 | GO:0002724 | regulation of T cell cytokine production | 1.7 | 3.00E-04 | **0.015** |
| 105 | GO:0022409 | positive regulation of cell-cell adhesion | 1.4 | 3.00E-04 | **0.015** |
| 106 | GO:0002287 | alpha-beta T cell activation involved in immune response | 1.6 | 3.00E-04 | **0.015** |
| 107 | GO:0002293 | alpha-beta T cell differentiation involved in immune response | 1.6 | 3.00E-04 | **0.015** |
| 108 | GO:0035745 | T-helper 2 cell cytokine production | 1.9 | 3.00E-04 | **0.015** |
| 109 | GO:0042249 | establishment of planar polarity of embryonic epithelium | 1.9 | 3.00E-04 | **0.015** |
| 110 | GO:0007259 | receptor signaling pathway via JAK-STAT | 1.5 | 3.00E-04 | **0.015** |

| | | | | | |
|---|---|---|---|---|---|
| 111 | GO:0003401 | axis elongation | 1.8 | 3.00E-04 | **0.016** |
| 112 | GO:0002696 | positive regulation of leukocyte activation | 1.3 | 3.00E-04 | **0.017** |
| 113 | GO:0043331 | response to dsRNA | 1.7 | 3.00E-04 | **0.017** |
| 114 | GO:0002430 | complement receptor mediated signaling pathway | 1.9 | 3.00E-04 | **0.017** |
| 115 | GO:0042035 | regulation of cytokine biosynthetic process | 1.5 | 3.00E-04 | **0.017** |
| 116 | GO:0042542 | response to hydrogen peroxide | 1.4 | 4.00E-04 | **0.017** |
| 117 | GO:0051250 | negative regulation of lymphocyte activation | 1.4 | 4.00E-04 | **0.018** |
| 118 | GO:0071692 | protein localization to extracellular region | 1.3 | 4.00E-04 | **0.018** |
| 119 | GO:0006935 | chemotaxis | 1.2 | 4.00E-04 | **0.018** |
| 120 | GO:0002791 | regulation of peptide secretion | 1.3 | 4.00E-04 | **0.018** |
| 121 | GO:0032642 | regulation of chemokine production | 1.6 | 4.00E-04 | **0.018** |
| 122 | GO:0045624 | positive regulation of T-helper cell differentiation | 1.8 | 4.00E-04 | **0.019** |
| 123 | GO:0032613 | interleukin-10 production | 1.6 | 4.00E-04 | **0.020** |
| 124 | GO:2000551 | regulation of T-helper 2 cell cytokine production | 1.9 | 4.00E-04 | **0.020** |
| 125 | GO:0018212 | peptidyl-tyrosine modification | 1.3 | 5.00E-04 | **0.020** |
| 126 | GO:0055010 | ventricular cardiac muscle tissue morphogenesis | 1.7 | 5.00E-04 | **0.020** |
| 127 | GO:0002830 | positive regulation of type 2 immune response | 1.8 | 5.00E-04 | **0.021** |
| 128 | GO:0034341 | response to interferon-gamma | 1.4 | 5.00E-04 | **0.021** |
| 129 | GO:0042088 | T-helper 1 type immune response | 1.6 | 5.00E-04 | **0.021** |
| 130 | GO:0006836 | neurotransmitter transport | 1.3 | 5.00E-04 | **0.021** |
| 131 | GO:0031294 | lymphocyte costimulation | 1.6 | 5.00E-04 | **0.021** |
| 132 | GO:0032602 | chemokine production | 1.5 | 5.00E-04 | **0.021** |
| 133 | GO:0010518 | positive regulation of phospholipase activity | 1.6 | 5.00E-04 | **0.021** |
| 134 | GO:0002833 | positive regulation of response to biotic stimulus | 1.3 | 5.00E-04 | **0.021** |
| 135 | GO:0090177 | establishment of planar polarity involved in neural tube closure | 1.9 | 5.00E-04 | **0.021** |
| 136 | GO:0090178 | regulation of establishment of planar polarity involved in neural tube closure | 1.9 | 5.00E-04 | **0.021** |
| 137 | GO:0043330 | response to exogenous dsRNA | 1.7 | 5.00E-04 | **0.021** |
| 138 | GO:1990266 | neutrophil migration | 1.5 | 5.00E-04 | **0.021** |
| 139 | GO:0000302 | response to reactive oxygen species | 1.3 | 5.00E-04 | **0.022** |
| 140 | GO:0050731 | positive regulation of peptidyl-tyrosine phosphorylation | 1.4 | 6.00E-04 | **0.023** |
| 141 | GO:0002573 | myeloid leukocyte differentiation | 1.4 | 6.00E-04 | **0.023** |
| 142 | GO:1904407 | positive regulation of nitric oxide metabolic process | 1.7 | 6.00E-04 | **0.023** |
| 143 | GO:0050870 | positive regulation of T cell activation | 1.3 | 6.00E-04 | **0.024** |
| 144 | GO:0042107 | cytokine metabolic process | 1.5 | 6.00E-04 | **0.024** |
| 145 | GO:0036005 | response to macrophage colony-stimulating factor | 1.9 | 6.00E-04 | **0.024** |
| 146 | GO:0036006 | cellular response to macrophage colony-stimulating factor stimulus | 1.9 | 6.00E-04 | **0.024** |
| 147 | GO:0032653 | regulation of interleukin-10 production | 1.6 | 6.00E-04 | **0.024** |
| 148 | GO:0002702 | positive regulation of production of molecular mediator of immune response | 1.5 | 6.00E-04 | **0.024** |
| 149 | GO:0032634 | interleukin-5 production | 1.9 | 6.00E-04 | **0.024** |
| 150 | GO:0032674 | regulation of interleukin-5 production | 1.9 | 6.00E-04 | **0.024** |

| | | | | | |
|---|---|---|---|---|---|
| 151 | GO:0050864 | regulation of B cell activation | 1.4 | 6.00E-04 | **0.024** |
| 152 | GO:0046637 | regulation of alpha-beta T cell differentiation | 1.6 | 7.00E-04 | **0.025** |
| 153 | GO:0050869 | negative regulation of B cell activation | 1.7 | 7.00E-04 | **0.027** |
| 154 | GO:0032722 | positive regulation of chemokine production | 1.6 | 8.00E-04 | **0.027** |
| 155 | GO:0002521 | leukocyte differentiation | 1.2 | 8.00E-04 | **0.027** |
| 156 | GO:0032943 | mononuclear cell proliferation | 1.3 | 8.00E-04 | **0.027** |
| 157 | GO:0051251 | positive regulation of lymphocyte activation | 1.3 | 8.00E-04 | **0.027** |
| 158 | GO:1904645 | response to amyloid-beta | 1.7 | 8.00E-04 | **0.027** |
| 159 | GO:0002822 | regulation of adaptive immune response based on somatic recombination of immune receptors built from immunoglobulin superfamily domains | 1.4 | 8.00E-04 | **0.029** |
| 160 | GO:0048660 | regulation of smooth muscle cell proliferation | 1.4 | 8.00E-04 | **0.029** |
| 161 | GO:0048659 | smooth muscle cell proliferation | 1.4 | 8.00E-04 | **0.030** |
| 162 | GO:0042110 | T cell activation | 1.2 | 9.00E-04 | **0.030** |
| 163 | GO:0050868 | negative regulation of T cell activation | 1.4 | 9.00E-04 | **0.030** |
| 164 | GO:0070661 | leukocyte proliferation | 1.3 | 9.00E-04 | **0.030** |
| 165 | GO:0002790 | peptide secretion | 1.2 | 9.00E-04 | **0.030** |
| 166 | GO:0009306 | protein secretion | 1.2 | 9.00E-04 | **0.030** |
| 167 | GO:0035592 | establishment of protein localization to extracellular region | 1.2 | 9.00E-04 | **0.030** |
| 168 | GO:0042267 | natural killer cell mediated cytotoxicity | 1.5 | 9.00E-04 | **0.030** |
| 169 | GO:0002711 | positive regulation of T cell mediated immunity | 1.5 | 9.00E-04 | **0.030** |
| 170 | GO:0042269 | regulation of natural killer cell mediated cytotoxicity | 1.6 | 9.00E-04 | **0.030** |
| 171 | GO:0043371 | negative regulation of CD4-positive, alpha-beta T cell differentiation | 1.7 | 9.00E-04 | **0.030** |
| 172 | GO:0002228 | natural killer cell mediated immunity | 1.5 | 1.00E-03 | **0.031** |
| 173 | GO:0002700 | regulation of production of molecular mediator of immune response | 1.4 | 1.00E-03 | **0.031** |
| 174 | GO:0031343 | positive regulation of cell killing | 1.5 | 1.00E-03 | **0.031** |
| 175 | GO:0002438 | acute inflammatory response to antigenic stimulus | 1.8 | 1.00E-03 | **0.031** |
| 176 | GO:0002819 | regulation of adaptive immune response | 1.4 | 1.00E-03 | **0.031** |
| 177 | GO:0032623 | interleukin-2 production | 1.5 | 1.00E-03 | **0.031** |
| 178 | GO:0032651 | regulation of interleukin-1 beta production | 1.5 | 1.00E-03 | **0.032** |
| 179 | GO:0032731 | positive regulation of interleukin-1 beta production | 1.6 | 1.00E-03 | **0.032** |
| 180 | GO:0046632 | alpha-beta T cell differentiation | 1.4 | 1.00E-03 | **0.032** |
| 181 | GO:0045217 | cell-cell junction maintenance | 1.8 | 1.00E-03 | **0.032** |
| 182 | GO:0018108 | peptidyl-tyrosine phosphorylation | 1.3 | 1.10E-03 | **0.033** |
| 183 | GO:0070664 | negative regulation of leukocyte proliferation | 1.5 | 1.10E-03 | **0.033** |
| 184 | GO:0010035 | response to inorganic substance | 1.2 | 1.10E-03 | **0.033** |
| 185 | GO:0046651 | lymphocyte proliferation | 1.3 | 1.10E-03 | **0.033** |
| 186 | GO:2000319 | regulation of T-helper 17 cell differentiation | 1.7 | 1.10E-03 | **0.034** |
| 187 | GO:0050866 | negative regulation of cell activation | 1.3 | 1.20E-03 | **0.036** |
| 188 | GO:0002673 | regulation of acute inflammatory response | 1.6 | 1.20E-03 | **0.036** |
| 189 | GO:0043122 | regulation of I-kappaB kinase/NF-kappaB signaling | 1.3 | 1.20E-03 | **0.036** |

| | | | | | |
|---|---|---|---|---|---|
| 190 | GO:0061081 | positive regulation of myeloid leukocyte cytokine production involved in immune response | 1.7 | 1.20E-03 | **0.036** |
| 191 | GO:0032729 | positive regulation of interferon-gamma production | 1.5 | 1.20E-03 | **0.036** |
| 192 | GO:0032740 | positive regulation of interleukin-17 production | 1.8 | 1.20E-03 | **0.036** |
| 193 | GO:0032732 | positive regulation of interleukin-1 production | 1.5 | 1.30E-03 | **0.037** |
| 194 | GO:0050900 | leukocyte migration | 1.2 | 1.30E-03 | **0.037** |
| 195 | GO:0051047 | positive regulation of secretion | 1.2 | 1.30E-03 | **0.038** |
| 196 | GO:0002431 | Fc receptor mediated stimulatory signaling pathway | 1.4 | 1.40E-03 | **0.040** |
| 197 | GO:0046634 | regulation of alpha-beta T cell activation | 1.4 | 1.40E-03 | **0.040** |
| 198 | GO:0007269 | neurotransmitter secretion | 1.4 | 1.40E-03 | **0.040** |
| 199 | GO:0099643 | signal release from synapse | 1.4 | 1.40E-03 | **0.040** |
| 200 | GO:0030889 | negative regulation of B cell proliferation | 1.7 | 1.50E-03 | **0.042** |
| 201 | GO:0001912 | positive regulation of leukocyte mediated cytotoxicity | 1.5 | 1.50E-03 | **0.042** |
| 202 | GO:0002715 | regulation of natural killer cell mediated immunity | 1.6 | 1.50E-03 | **0.042** |
| 203 | GO:0030888 | regulation of B cell proliferation | 1.5 | 1.50E-03 | **0.042** |
| 204 | GO:0043405 | regulation of MAP kinase activity | 1.3 | 1.60E-03 | **0.043** |
| 205 | GO:0045429 | positive regulation of nitric oxide biosynthetic process | 1.6 | 1.60E-03 | **0.044** |
| 206 | GO:0050885 | neuromuscular process controlling balance | 1.6 | 1.60E-03 | **0.044** |
| 207 | GO:2000379 | positive regulation of reactive oxygen species metabolic process | 1.5 | 1.60E-03 | **0.044** |
| 208 | GO:0060026 | convergent extension | 1.8 | 1.60E-03 | **0.044** |
| 209 | GO:0033674 | positive regulation of kinase activity | 1.2 | 1.70E-03 | **0.044** |
| 210 | GO:0038094 | Fc-gamma receptor signaling pathway | 1.4 | 1.70E-03 | **0.044** |
| 211 | GO:0050829 | defense response to Gram-negative bacterium | 1.6 | 1.70E-03 | **0.045** |
| 212 | GO:0050730 | regulation of peptidyl-tyrosine phosphorylation | 1.3 | 1.70E-03 | **0.045** |
| 213 | GO:1903426 | regulation of reactive oxygen species biosynthetic process | 1.5 | 1.70E-03 | **0.045** |
| 214 | GO:1901224 | positive regulation of NIK/NF-kappaB signaling | 1.4 | 1.70E-03 | **0.045** |
| 215 | GO:2000516 | positive regulation of CD4-positive, alpha-beta T cell activation | 1.6 | 1.70E-03 | **0.045** |
| 216 | GO:0002709 | regulation of T cell mediated immunity | 1.5 | 1.80E-03 | **0.045** |
| 217 | GO:0002824 | positive regulation of adaptive immune response based on somatic recombination of immune receptors built from immunoglobulin superfamily domains | 1.4 | 1.80E-03 | **0.045** |
| 218 | GO:0060740 | prostate gland epithelium morphogenesis | 1.8 | 1.80E-03 | **0.045** |
| 219 | GO:0097191 | extrinsic apoptotic signaling pathway | 1.3 | 1.80E-03 | **0.045** |
| 220 | GO:0030101 | natural killer cell activation | 1.5 | 1.80E-03 | **0.045** |
| 221 | GO:0045619 | regulation of lymphocyte differentiation | 1.3 | 1.80E-03 | **0.045** |
| 222 | GO:1904892 | regulation of receptor signaling pathway via STAT | 1.4 | 1.80E-03 | **0.045** |
| 223 | GO:0006450 | regulation of translational fidelity | 1.7 | 1.90E-03 | **0.047** |
| 224 | GO:0034614 | cellular response to reactive oxygen species | 1.3 | 1.90E-03 | **0.048** |

[a] NES = normalized enrichment score.
[b] p = two sided p-value.
[c] q = two-sided false discovery rate (FDR) Benjamini-Hochberg q-value.

**Supplemental Table 5.** Differentially expressed host genes in relation to HIV Intact DNA in the Total Study Population (top panel) and the European ancestry subgroup (bottom panel), at a Benjamini-Hochberg false discovery rate (FDR) of q<0.25. Two-fold higher level of HIV intact DNA was associated with upregulation of genes involved in glycogen degradation (*AGL*) and inhibits thrombus (clot) degradation (*PLGLB1*).

| HIV Intact DNA | | | | | | |
|---|---|---|---|---|---|---|
| **Gene** | **Gene Name** | **p[a]** | **q[b]** | **FC[c]** | **% Change[d]** | **Description** |
| **Total Study Population** | | | | | | |
| NA | | | | | | |
| **European Ancestry Subgroup** | | | | | | |
| *AGL* | amylo-alpha-1, 6-glucosidase, 4-alpha-glucanotransferase | 2.09E-05 | 0.23 | 1.009 | 0.9 | *AGL* (AGL, Amylo-Alpha-1, 6-Glucosidase, 4-Alpha-Glucanotransferase) gene is involved in glycogen metabolism (process of breaking down stored glucose into glucose molecules for immediate glucose release and availability) [148-150]. |
| *PLGLB1* | plasminogen like B1 | 2.39E-05 | 0.23 | 1.060 | 6.0 | *PLGLB1 (*plasminogen like B1) gene is involved in thrombin clot degradation) [151-153]. PLGLB1 has previously been linked to a clonally expanded HIV-1 provirus (integrated in the opposite direction) from a patient with squamous cell carcinoma [155]. |

[a] p = two sided p-value.
[b] q = two-sided false discovery rate (FDR) Benjamini-Hochberg q-value. Bold font denotes genes with q<0.05.
[c] FC = fold-change in host gene expression per two-fold change in copies of HIV from multivariate model adjusted for age, sex, nadir CD4+ T cell count, timing of ART initiation, ancestry (PCs), and residual variability (probabilistic estimation of expression residuals, PEERs).
[d] % Change = percent change in host gene expression per two-fold change in copies of HIV.

**Supplemental Table 6.** Gene set enrichment analyses (GSEA) of ranked differentially expressed genes in relation to HIV intact DNA using the Gene Ontology Biological Processes (GO-BP) database. Genes sets with Benjamini-Hochberg false discovery rate (FDR)-adjusted q<0.25 are shown for the total study population (top panel) and for the European ancestry subgroup (bottom panel). Gene sets where q<0.05 are shown in bold font.

| HIV Intact DNA | | | | | |
|---|---|---|---|---|---|
| | **GO ID** | **Description** | **NES[a]** | **p[b]** | **q[c]** |
| **Total Study Population** | | | | | |
| NA | | | | | |
| **European Ancestry Subgroup** | | | | | |
| 1 | GO:0043312 | neutrophil degranulation | 1.3 | 1.49E-05 | **0.046** |
| 2 | GO:0002283 | neutrophil activation involved in immune response | 1.3 | 1.91E-05 | **0.046** |
| 3 | GO:0043299 | leukocyte degranulation | 1.3 | 2.47E-05 | **0.046** |
| 4 | GO:0002444 | myeloid leukocyte mediated immunity | 1.3 | 4.13E-05 | 0.058 |
| 5 | GO:0002275 | myeloid cell activation involved in immune response | 1.3 | 6.15E-05 | 0.060 |
| 6 | GO:0002446 | neutrophil mediated immunity | 1.3 | 6.49E-05 | 0.060 |
| 7 | GO:0042119 | neutrophil activation | 1.3 | 7.50E-05 | 0.060 |
| 8 | GO:0036230 | granulocyte activation | 1.3 | 1.00E-04 | 0.085 |
| 9 | GO:0019377 | glycolipid catabolic process | 1.9 | 3.00E-04 | 0.157 |
| 10 | GO:0016311 | dephosphorylation | 1.3 | 3.00E-04 | 0.162 |
| 11 | GO:0046479 | glycosphingolipid catabolic process | 1.9 | 4.00E-04 | 0.205 |
| 12 | GO:0060037 | pharyngeal system development | 1.9 | 5.00E-04 | 0.224 |

[a] NES = normalized enrichment score.
[b] p = two sided p-value.
[c] q = two-sided false discovery rate (FDR) Benjamini-Hochberg q-value.

**Supplemental Figure 2.** Proposed model for the observed inverse associations between HIV total DNA and tumor suppressor gene (*NBL1, P3H3*) expression (q<0.05). Given the known function of these genes, we hypothesize that these tumor suppressor genes might act as "checks" on cell proliferation, thereby restricting the size of the total HIV DNA reservoir.

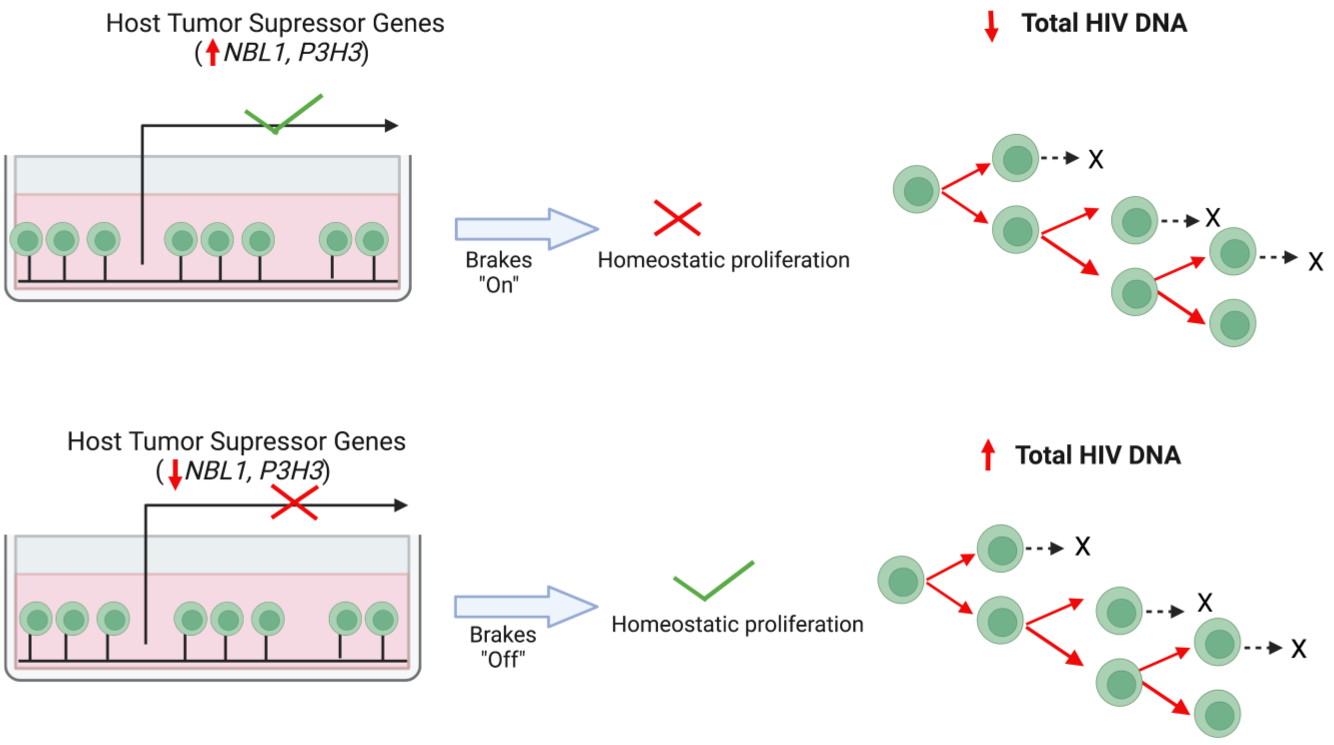

**Supplemental Figure 3.** Proposed model for the observed inverse associations between HIV unspliced RNA and several host genes involved in innate immune activation and inflammation (q<0.05). Given the known function of these genes, e.g., cytokine production (*IL1A, CSF3, TNFAIP6*), chemokine signaling (*CXCL3, CXCL10*), and pathogen pattern recognition (*TLR7*), we hypothesize that individuals with a more "transcriptionally active" reservoir may downregulate host genes perpetuating immune activation and inflammation (a). For the observed inverse association with *KCNJ2*, a gene encoding for an inwardly rectifying potassium channel (Kir2.1), previously shown to enhance HIV entry and viral release [32], we hypothesize that individuals with a more "transcriptionally active" reservoir may suppress host gene expression that might further promote HIV infectivity (b).

(a)

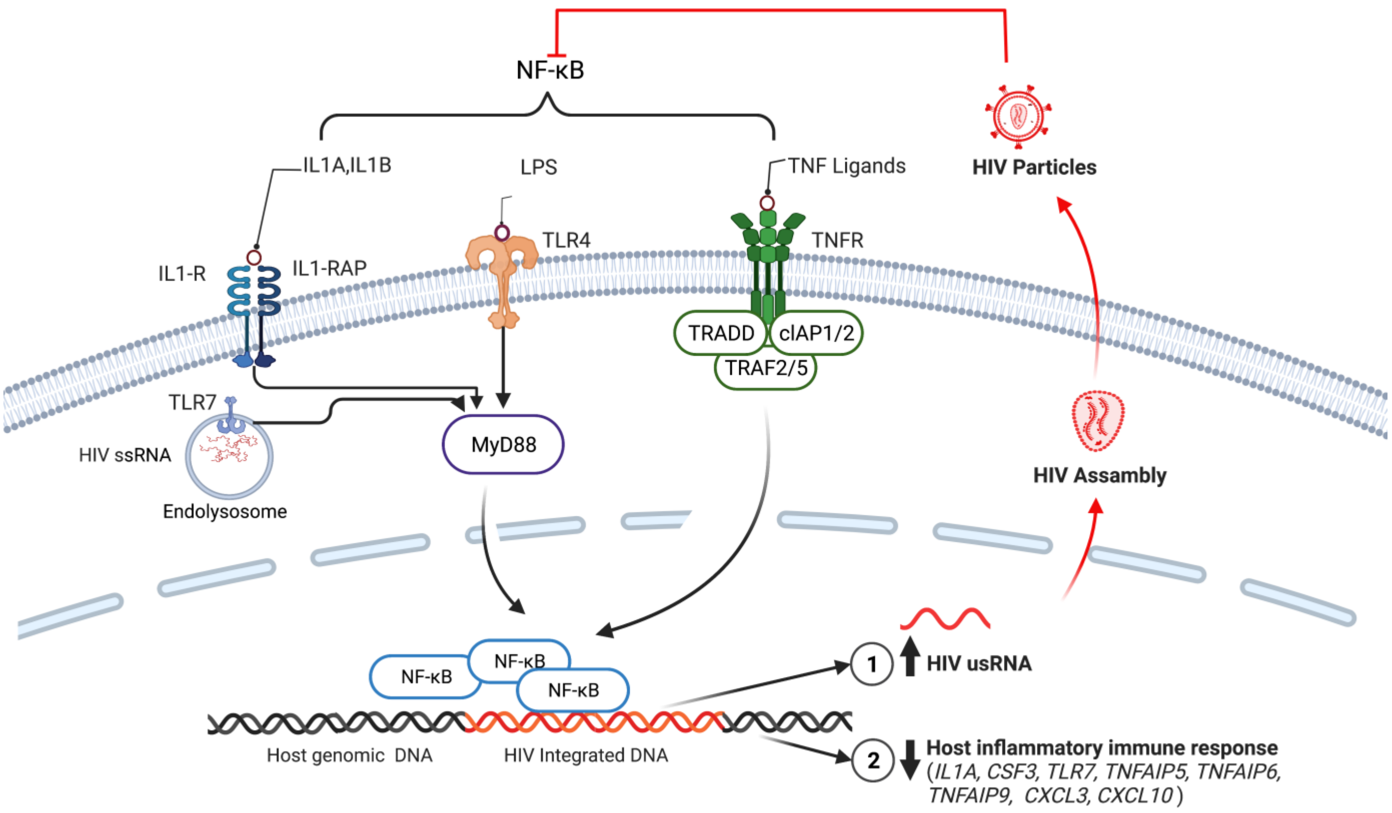

(b)

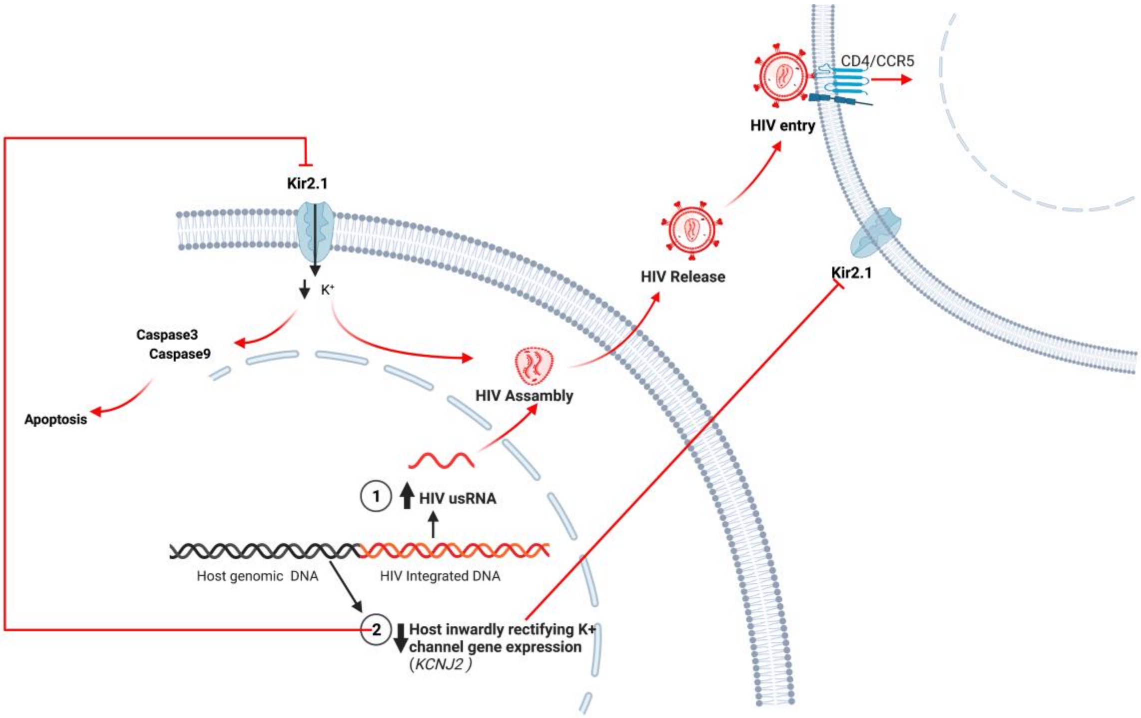
CD4/CCR5
HIV entry
Kir2.1
K+
HIV Release
Kir2.1
Caspase3
Caspase9
HIV Assambly
Apoptosis
1
HIV usRNA
Host genomic DNA
HIV Integrated DNA
2
Host inwardly rectifying K+
channel gene expression
(KCNJ2)

**Supplemental Figure 4.** HIV intact DNA was undetectable in 48% of our measured samples. Proposed model for the observed positive trend between HIV intact DNA and two genes, *PLGLB1* and *AGL* in the European ancestry subgroup (q<0.25). Given the known function of these genes, these trends might suggest that upregulation of *PLGLB1* (encodes for a protein that inhibits thrombus degradation) and *AGL* (encodes for an enzyme which is involved in glycogen degradation) reflect an association between the "replication-competent" HIV intact DNA reservoir and host inflammatory sequelae.

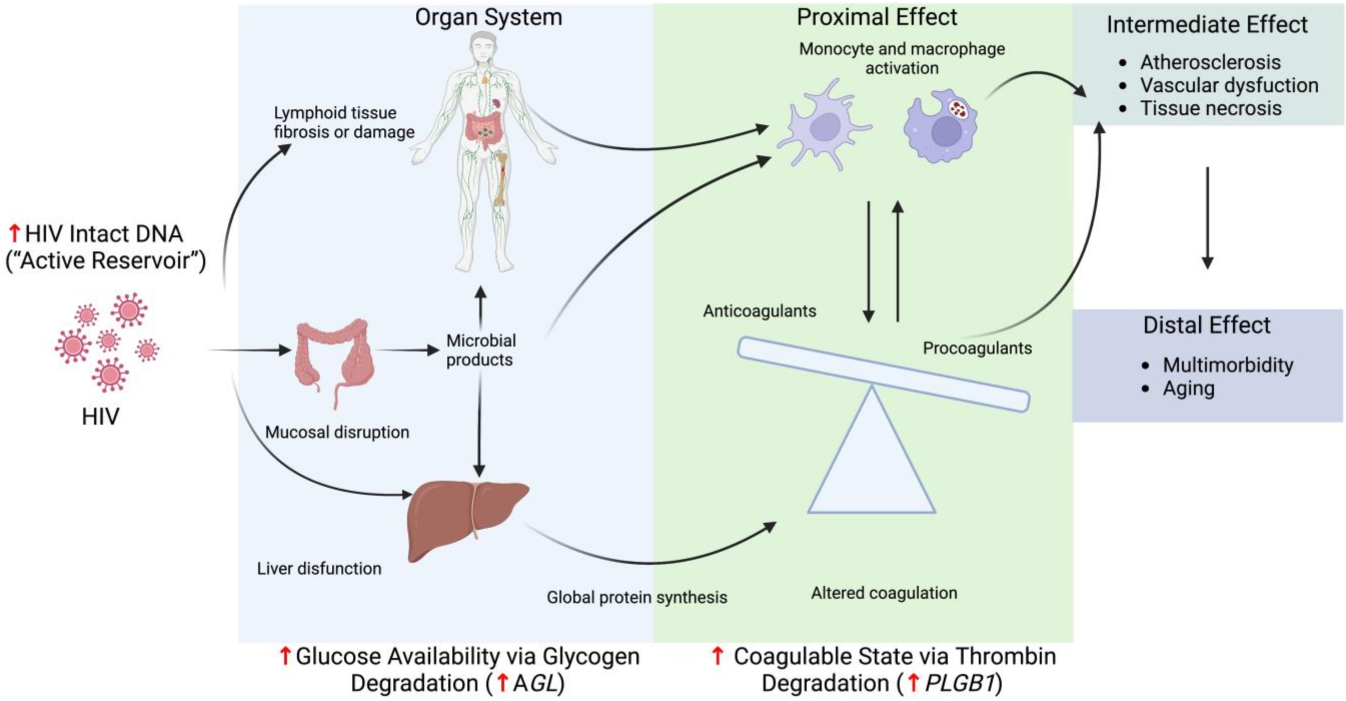
